# MEA-NAP compares microscale functional connectivity, topology, and network dynamics in organoid or monolayer neuronal cultures

**DOI:** 10.1101/2024.02.05.578738

**Authors:** Timothy PH Sit, Rachael C Feord, Alexander WE Dunn, Jeremi Chabros, David Oluigbo, Hugo H Smith, Lance Burn, Elise Chang, Alessio Boschi, Yin Yuan, George M Gibbons, Mahsa Khayat-Khoei, Francesco De Angelis, Erik Hemberg, Martin Hemberg, Madeline A Lancaster, Andras Lakatos, Stephen J Eglen, Ole Paulsen, Susanna B Mierau

**Author notes:** equal contribution (Timothy PH Sit, Rachael C Feord).

## Abstract

Microelectrode array (MEA) recordings are commonly used to compare firing and burst rates in neuronal cultures. MEA recordings can also reveal microscale functional connectivity, topology, and network dynamics—patterns seen in brain networks across spatial scales. Network topology is frequently characterized in neuroimaging with graph theoretical metrics. However, few computational tools exist for analyzing microscale functional brain networks from MEA recordings. Here, we present a MATLAB MEA network analysis pipeline (MEA-NAP) for raw voltage time-series acquired from single- or multi-well MEAs. Applications to 3D human cerebral organoids or 2D human-derived or murine cultures reveal differences in network development, including topology, node cartography, and dimensionality. MEA-NAP incorporates multi-unit template-based spike detection, probabilistic thresholding for determining significant functional connections, and normalization techniques for comparing networks. MEA-NAP can identify network-level effects of pharmacologic perturbation and/or disease-causing mutations and, thus, can provide a translational platform for revealing mechanistic insights and screening new therapeutic approaches.

## Introduction

Microelectrode array (MEA) recordings from in vitro models of brain development and disease offer a cellular-scale platform for mechanistic and therapeutic studies in 2D murine or human-derived neuronal cultures and lately in 3D human cortical organoids. The parallel streams of information acquired from the MEA enable investigation of the patterns of functional connectivity and network topology that develop in these microscale functional networks in vitro ^1,2^. These features provide an electrophysiological phenotype of the effects of disease-causing genetic mutations on the efficiency of information processing ^3^ and can be used to understand differences in computational performance in human cerebral organoids ^4^.

Patterns of functional connectivity and network topology observed at whole-brain level have revealed multiple organizing principles that are correlated with higher efficiency of cognitive processes, including the development of hubs and small-world topology ^5^. Computational tools for analyzing networks at the whole-brain level are widely used, including the Brain Connectivity Toolbox ^6^. Similar patterns and motifs in the functional connectivity and network topology have been observed at the microscale level in murine monolayer ^7,8^ and human-derived organoid ^9^ neuronal cultures. Yet relatively few studies of MEA recordings from 2D or 3D neuronal cultures, or complex organoids, go beyond measures of activity or correlation alone ^10^. This is primarily due to the limited availability of computational tools to reveal significant functional connectivity and to compare network metrics between cultures at different ages, species, cell-type diversity, or in different conditions that affect network size and/or density ^10^.

Here, we created the MEA network analysis pipeline (MEA-NAP) as a diagnostic tool for exploring functional connectivity and network topology in MEA recordings. Our work introduces several network analysis methodologies to facilitate analyses of microscale functional neuronal networks. We demonstrate their utility by analyzing several datasets from human-derived and murine cultures. Together, these methods expand our understanding of brain network development and dysfunction, and they make it possible to use the methods popularized by the Brain Connectivity Toolbox across spatial scales. MEA-NAP is suitable for users without prior knowledge of network neuroscience, since it provides a single toolbox for batch analysis of entire experiments of MEA recordings. This streamlines the analysis of MEA recordings, combining robust spike (action potential) detection using template-based methods, comparison of action potential firing and burst rates, inferring functional connectivity from significant correlated activity, and comparison of network features including the topology and network dynamics. Importantly, MEA-NAP provides advances over existing tools through integrated validation, visualization, and statistical tools. In particular, MEA-NAP provides multi-unit spike detection with validation tools for comparing different template- and threshold-based methods. MEA-NAP automatically generates figures with scaling to individual MEA recordings and to the entire dataset to facilitate comparisons at the electrode, recording, and entire-dataset level. MEA-NAP successfully identifies network features in the development of 2D or 3D, murine or human-derived neuronal cultures recorded with Multi-channel Systems (MCS) standard density single-well (60 electrodes) or Axion Biosystems multi-well (64 electrodes) MEA systems. MEA-NAP compares patterns of network activity over different time points (e.g., development) and between conditions (e.g., genetic mutations, drug application, other perturbation experiments) revealing how functional networks form in different types of murine and human-derived neuronal cultures and how they are affected by pharmacologic or optogenetic stimulation.

## Results

### MEA-NAP is a user-friendly MATLAB tool

MEA-NAP is designed to batch analyze MEA recordings from an entire experiment for users who have little-to-no expertise in MATLAB, or network science, and to be customizable for more experienced users. The inputs to the pipeline are MEA recordings (raw or filtered voltage time series) imported to MATLAB. Figure 1 provides an overview of MEA-NAP’s five steps. In Step 1, spike detection is performed to identify action potentials detected from the multi-unit activity at individual electrodes (nodes). In Step 2, the neuronal activity, including mean firing and network burst rates, are compared by age and group (e.g., genotype). In Step 3, significant functional connections (edges) between pairs of nodes are inferred from correlated spiking activity using the spike time tiling coefficient (STTC) ^11^ and probabilistic thresholding. In Step 4, graph theoretical metrics from the Brain Connectivity Toolbox and other network metrics are evaluated for each age and experimental condition. In Step 5, statistical comparisons and feature selection are performed. At each step, MEA-NAP automatically generates informative figures to facilitate visualization of neuronal activity and network features at the electrode, recording, and entire-dataset level and validation of spike detection, probabilistic thresholding, and other parameters. Therefore, MEA-NAP provides a diagnostic tool for examining microscale functional network features in neuronal cultures. We exploited this approach to robustly apply network analysis methods from the wider field of network neuroscience to MEA recordings of neuronal cultures to facilitate the comparison of microscale functional networks in early development and disease in in vitro human-derived or animal, 2D or 3D models.

**Figure 1.**
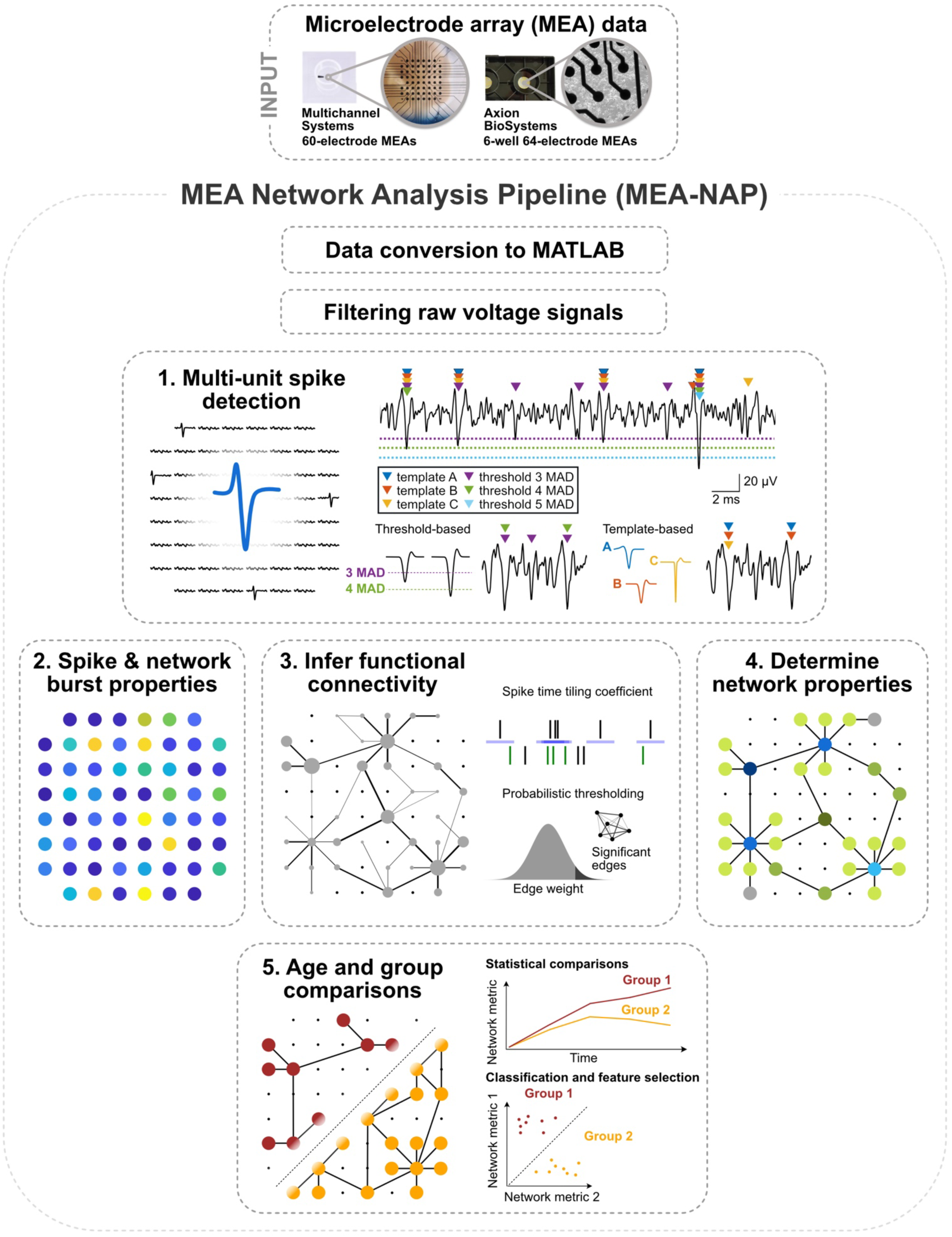
Overview of the MEA-NAP. Illustration of microelectrode array (MEA) network analysis pipeline (MEA-NAP) steps 1-5. MEA data from 60- (Multichannel systems single-well) or 64-electrode (Axion Biosystems, 6-well plates) MEA systems serves as input to the pipeline. The data is converted to MATLAB format, and the raw voltage signal is filtered. **1. Multi-unit spike detection** is performed to extract the action potential time series from each electrode. Left, action potentials detected from multiple electrodes in the MEA. Right top, sample voltage trace from a single electrode showing spikes detected (colored arrows) by different template-based and threshold-based methods. Right bottom, example action potentials detected by median absolute deviation (MAD) 3 or 4 thresholds. Example wavelets (A-C) serve as templates for detecting spikes. **2. Neuronal activity** compares the spike and network burst properties within and between MEA recordings. **3. Functional connectivity** is determined from significant pairwise correlation of the neuronal activity (left) by combining the spike time tiling coefficient with probabilistic thresholding (right). **4. Network activity** compares multiple network topological and dynamic features. Illustration highlights node cartography for identify hub and non-hub nodal roles. **5. Statistical analysis** illustrates age and group comparisons and feature selection.

### Multi-unit template-based spike detection shows high sensitivity and specificity in a 3D human cerebral organoid

Human cerebral organoids offer an exciting platform for studying brain development in health and disease because they recapitulate both the cell-type diversity and some of the 3D architecture, such as cortical layering, of the human brain ^3^. We previously identified functional connectivity in a novel air-liquid interface cortical organoid (ALI-CO) model ^9^. We sought to expand and improve the tools for characterizing neuronal activity, functional connectivity, and network topology in ALI-COs and other 3D human cerebral organoids models through new features in MEA-NAP. One of the key challenges in analysis of MEA recordings from 3D human cerebral organoids (Figure 2A) is the detection of multi-unit activity due to the higher density of neurons near each electrode, compared to 2D cultures. The amplitude of the action potentials from different neurons near an individual electrode will vary based on the distance from the electrode and other features (Figure 2B), which can lead to a trade-off in sensitivity or specificity when threshold-based methods, such as those commonly included with the MEA system acquisition software, are used.

**Figure 2.**
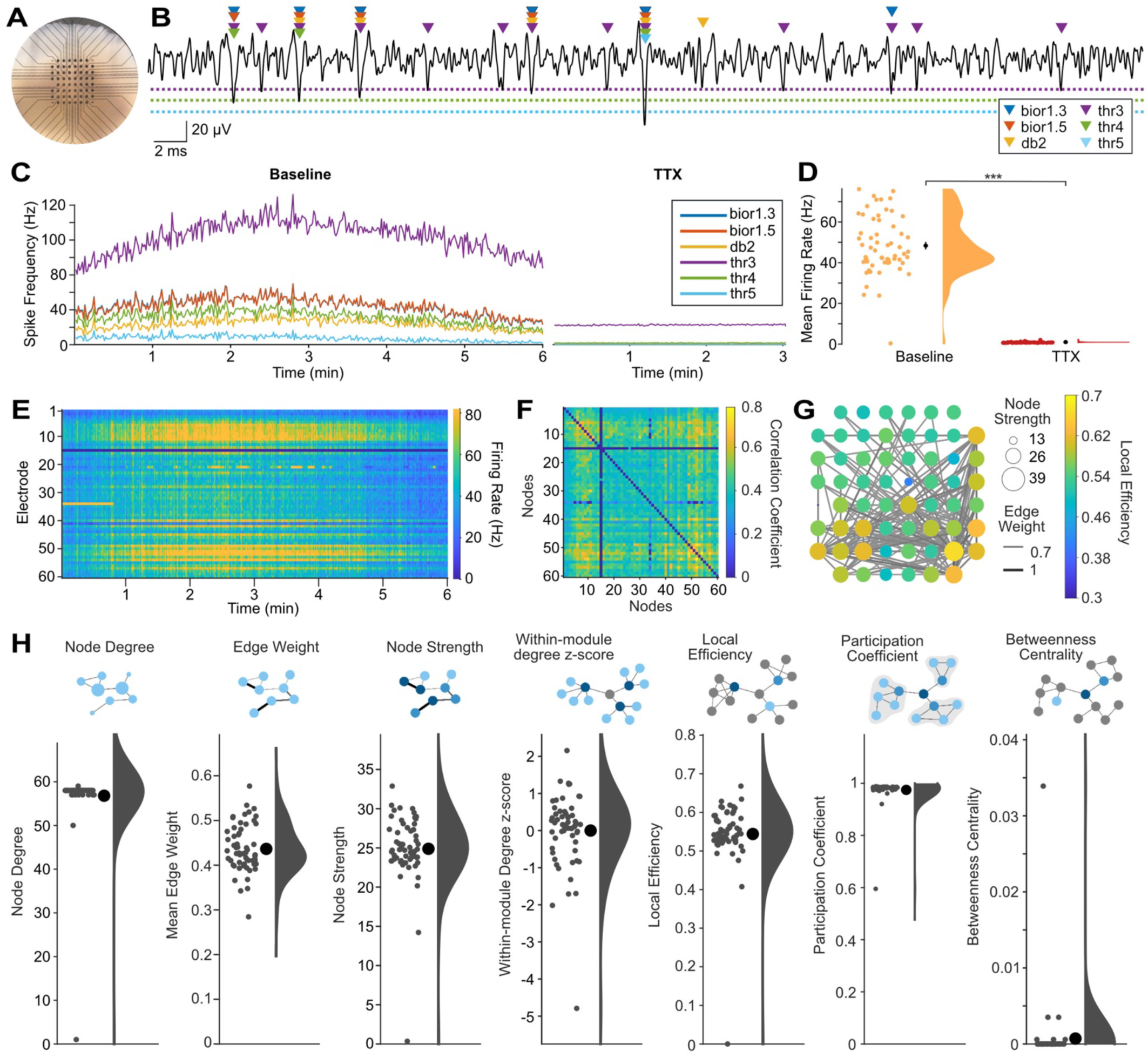
MEA-NAP offers template-based multi-unit spike detection and automated analysis of functional connectivity and network topology in 3D human cerebral organoids. **A.** Representative photo of an air-liquid interface cortical organoid (ALI-CO) on the microelectrode array (MEA). Scale: electrodes (black circles) are 200 μm apart. **B.** Sample 60-ms-long voltage trace (black line) from a single electrode showing comparison of spikes detected by the continuous wavelet transform with templates bior1.3, bior1.5, and db2 and by a median absolute deviation threshold of 3, 4 and 5 (colored triangles) in a days-in-vitro (DIV) 154 ALI-CO. Dashed lines show thresholds (3, 4, 5 in the same colors as triangles). Scale bars: vertical 20 uV, horizontal, 2 ms. **C.** Comparison of spike detection methods with running averages of spikes per second for all electrodes in the MEA for each method (colored lines, see legend) over a 6-minute baseline recording (left) and after application of 1 μM tetrodotoxin (TTX) to block action potentials (right). **D.** Scatter with half-violin plot of density curve for mean firing rate by electrode during baseline recording and after TTX using the multi-unit template-based spike detection. Mean (black circles) ± SEM (error bars). *** p<0.00001 (paired two-tailed t-test). Where error bars are not visible indicates the SEM was smaller than the size of the circle. **E.** Raster plot shows firing rate (spikes per second, color bar on right) for each electrode (row) over the 6-minute recording using the multi-unit template-based spike detection. **F.** Adjacency matrix shows correlation coefficient (spike time tiling coefficient, color bar) for significant functional connections (edges) between pairs of electrodes (nodes). Significant edges determined using probabilistic thresholding. **G.** Graph of network illustrating functional connectivity in the ALI-CO. Nodes show node strength (circle size) and edge weight (line thickness) strength of connectivity. Only edges with edge weight greater than 0.55 are plotted to show the strongest functional connections. Node color (scale bar) represents local efficiency. **H.** Summary of nodal-level graph metrics for the DIV154 ALI-CO. Diagrams depict nodal-level graph metrics. Scatter plots show values for individual nodes (small gray circles), mean ± SEM (large black circle with error bars), and density curves.

To address this challenge, MEA-NAP has the option to run one or multiple spike detection methods, and it provides validation tools to compare how the spike detection methods perform on specific experimental datasets (Figure 2C). In contrast to most publicly available and commercial tools for spike detection, MEA-NAP offers template-based spike detection with the continuous wavelet transform ^12,13^ to identify spikes based on their waveform, using MATLAB built-in wavelets. The main advantage of template-based over threshold-based spike detection methods is that they can increase both sensitivity and specificity by identifying action potentials based on their morphology. Template-based spike detection is less sensitive to large artifacts (noise deflections) and differences in spike frequency between recordings than threshold-based methods. We compared the template-based spike detection using three MATLAB built-in wavelets (bior1.5, bior1.3, db2) and threshold-based spike detection using three mean absolute deviations (3,4,5) on MEA recordings from a human cerebral organoid (Figure 2A) before and after application of 1 μM tetrodotoxin (TTX), which effectively inhibits action potentials. The wavelets detect more spikes than thresholds 4 and 5 (Figures 2B and 2C). Threshold 3 detects a larger number of spikes, as shown in the running average of the number of spikes detected by each method in the baseline recording (Figure 2C, left panel); however, the lower specificity of threshold 3 than the template-based methods is revealed in the TTX condition (Figure 2C, right panel). Multi-unit spike detection of action potentials detected by one or more wavelets was highly specific for action potentials (Figure 2D, p<1e-5, paired t-test) and was used to create a raster plot of the neuronal activity in the MEA over the 6-minute recording (Figure 2E). Importantly, depending on the signal-to-noise ratio and the goodness of fit of selected templates to the action potential morphology, different spike detection methods may perform better on some datasets (e.g., 2D versus 3D, human versus murine cultures) than others. In MEA-NAP, users can select a single method (wavelet- or threshold-based) or to combine methods to achieve multi-unit spike detection. When combined, spike times detected by different methods are compared first to ensure that spikes detected by multiple wavelets or thresholds are only counted once for the downstream comparisons of neuronal activity and network features.

To infer significant functional connectivity, MEA-NAP applies probabilistic thresholding to determine significant functional connections using the spike time tiling coefficient (STTC) ^11^. Probabilistic thresholding facilitates robust comparison of network features where the network size and density may vary—such as with age or disease condition—while retaining weak but significant connections ^14,15^. MEA-NAP calculates the STTC for the spike trains from each pair of electrodes. Next circular shifts are performed on one of the spike trains for each pair. This method of rearranging the spike train preserves the number of action potentials and the distribution of interspike intervals. The STTC is calculated again for each iteration for each pair. A functional connection was determined when the real STTC was greater than 95th percentile of the STTC values from the circular shifts. The significant pairwise correlations are represented in the adjacency matrix (Figure 2F), and the functional connectivity is visualized in the spatial arrangement of the MEA (Figure 2G). Graph theoretical metrics were calculated and examples of node-level features in the network are shown (Figure 2H). The MEA recording from the days-in-vitro (DIV) 154 ALI-CO shows dense network activity with high mean node degree (number of significant connections per node) and participation coefficient (Figure 2H). This indicates that the neurons in the ALI-CO have many connections distributed throughout the network and participate in the overall network activity, rather than their activity being restricted to smaller modules or subcommunities of neurons within the ALI-CO.

### MEA-NAP reveals development of functional connectivity and network topology in 2D human iPSC-derived NGN2 cortical cultures

2D human iPSC-derived neuronal cultures show a developmental increase in firing rates and bursting activity that may reveal disease-related phenotypes at the microscale ^16,17^. MEA-NAP efficiently analyzes MEA recordings from neurogenin 2 (NGN2) iPSC-derived cortical cultures (Figure 3A), quantifying and visualizing firing rates and bursting activity, including network bursts. Raster plots of individual recordings from the same culture illustrate the increase in firing rate from DIV14 to 35 and the development of network bursts by DIV 35 (Figure 3B). Network bursts were defined as bursts of action potentials detected in 3 or more electrodes within a given time frame, or lag, in milliseconds using the ISI_N_ method ^18^. To address the effect of age on the development of neuronal activity, functional connectivity, and network topology metrics, we applied a one-way ANOVA and the Tukey-Kramer method to adjust for multiple pairwise comparisons. NGN2 iPSC-derived cortical cultures (n=15) showed a significant increase in the number of active electrodes (p=8e-7), mean firing rate (p=7e-6), fraction of bursts occurring in network bursts (p=7e-4), mean network burst rate (p=0.02) between DIV 14-35 (Figures 3C-3F). This was accompanied by a significant decrease in the interspike interval (ISI) within (p=2e-4) and outside network bursts (p=4e-4; Figures 3G and 3H). MEA-NAP goes beyond existing computational tools to perform batch analysis comparing the development of functional connectivity and network topology (Table 1). MEA-NAP generated graphs of the significant functional connectivity in MEA recordings from individual cultures at DIV 14-35 (Figure 3I). As illustrated by the size of the node, the node degree (number of significant connections) was not uniform across all electrodes, with the neurons near a subset of electrodes being more connected. The edge weight (strength of connectivity) also varied. The network density (proportion of significant connections from all possible connections, p=0.002) and top 10% of edge weight (p=4e-5) significantly increased from DIV 14-35 (Figures 3J and 3K). The number of modules significantly decreased from DIV 14-35 (p=0.01), while the mean participation coefficient (proportion of a node’s connections distributed among different modules) in the top 10% of nodes significantly increased from DIV 14-35 (p=0.001). At DIV 14-28, the network topology approaches a randomly connected network (Figure 3N). However, by DIV35, small-world networks form as well as networks which approach a lattice-like (more highly connected) network. The mean small-worldness coefficient (ω)^19^ approached zero, indicating small-world topology, at DIV35 compared to more positive values at DIV 14-28 (p=3e-4), indicating a more random network. MEA-NAP includes visualization tools for comparing the choice of STTC lag on downstream graph theoretical metrics (Figure 3O). The difference in small-worldness at DIV35 was not dependent on the choice of lag (10, 25 or 50 ms) used to determine significant functional connections with the STTC.

**Table 1.**
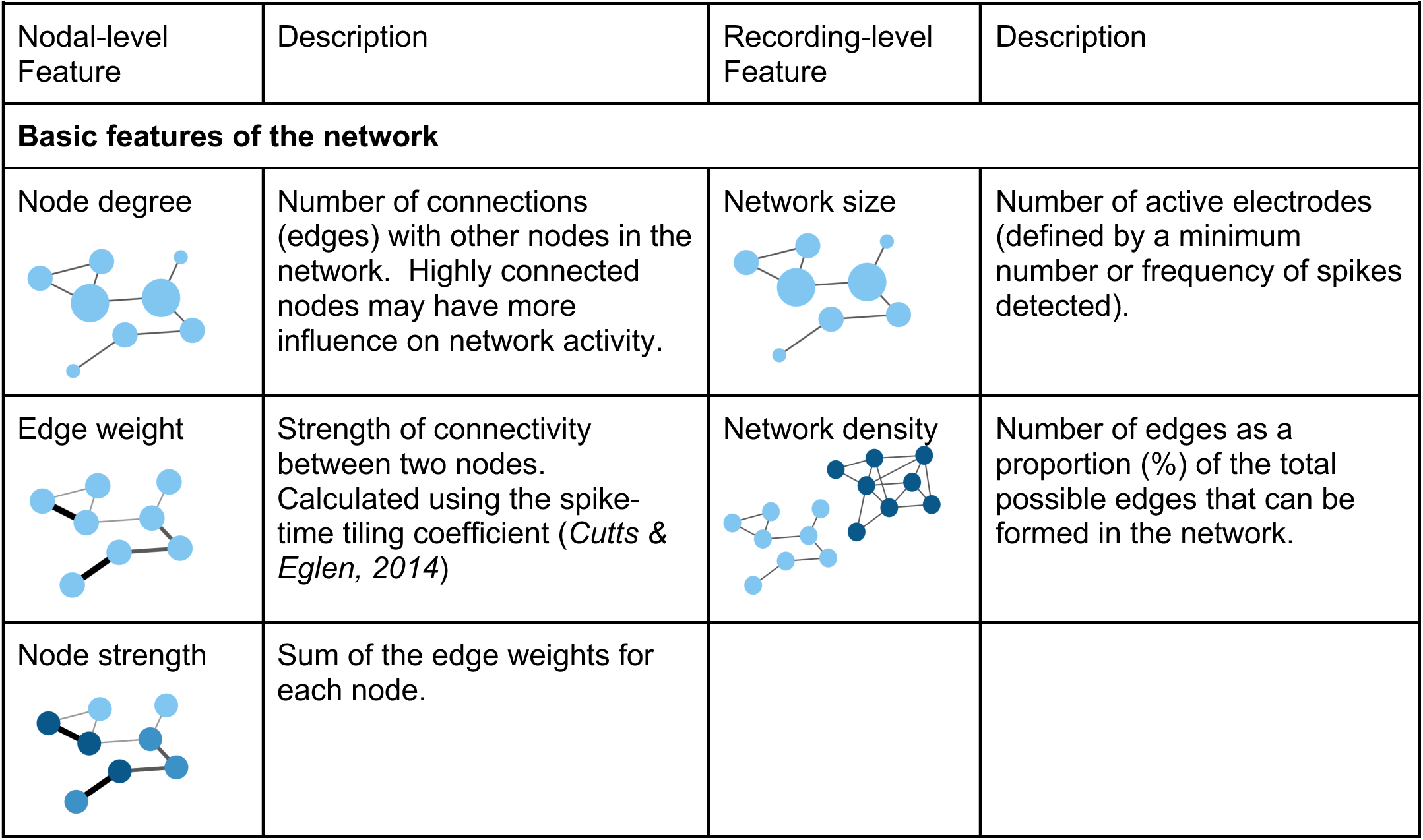
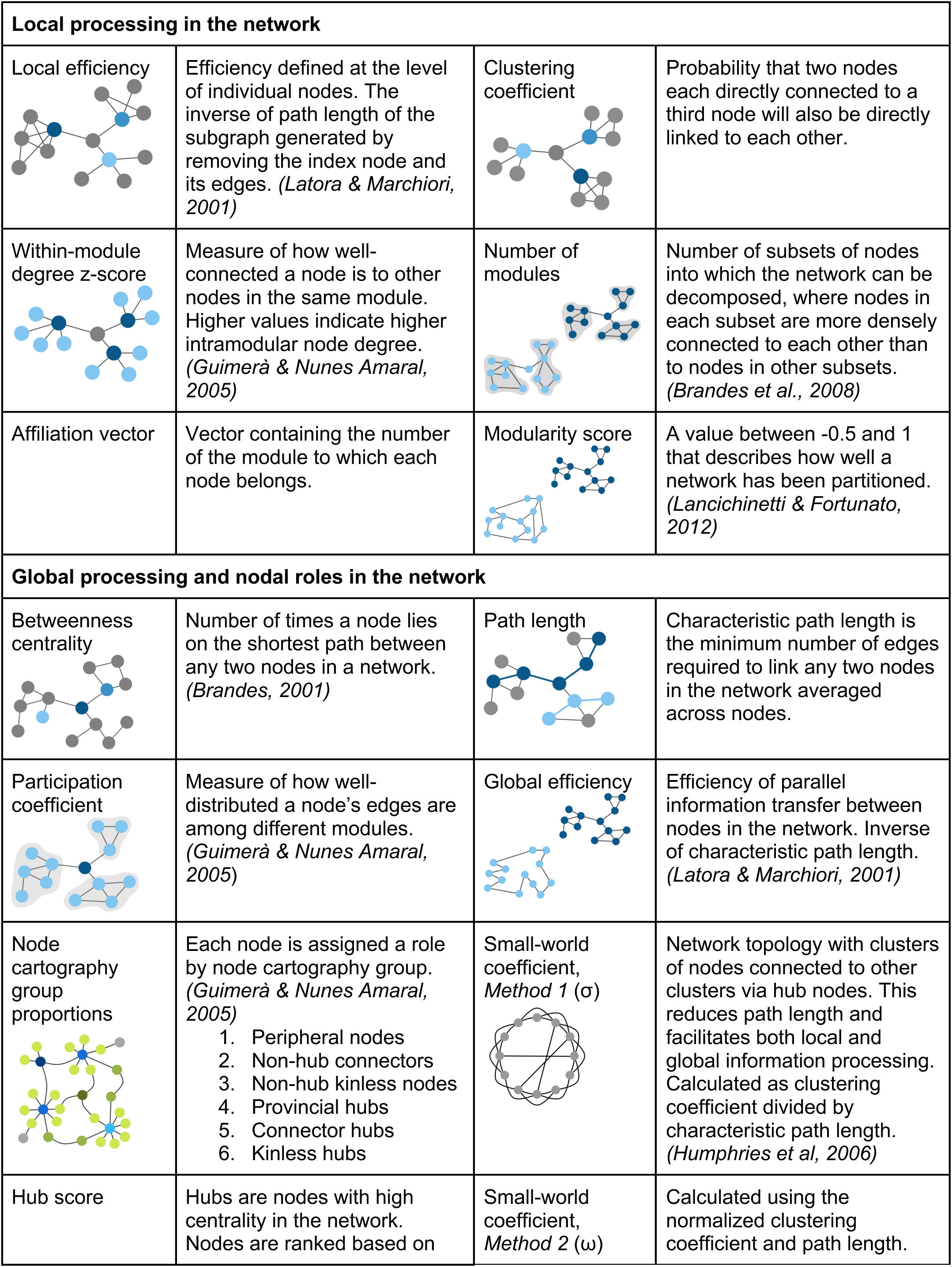
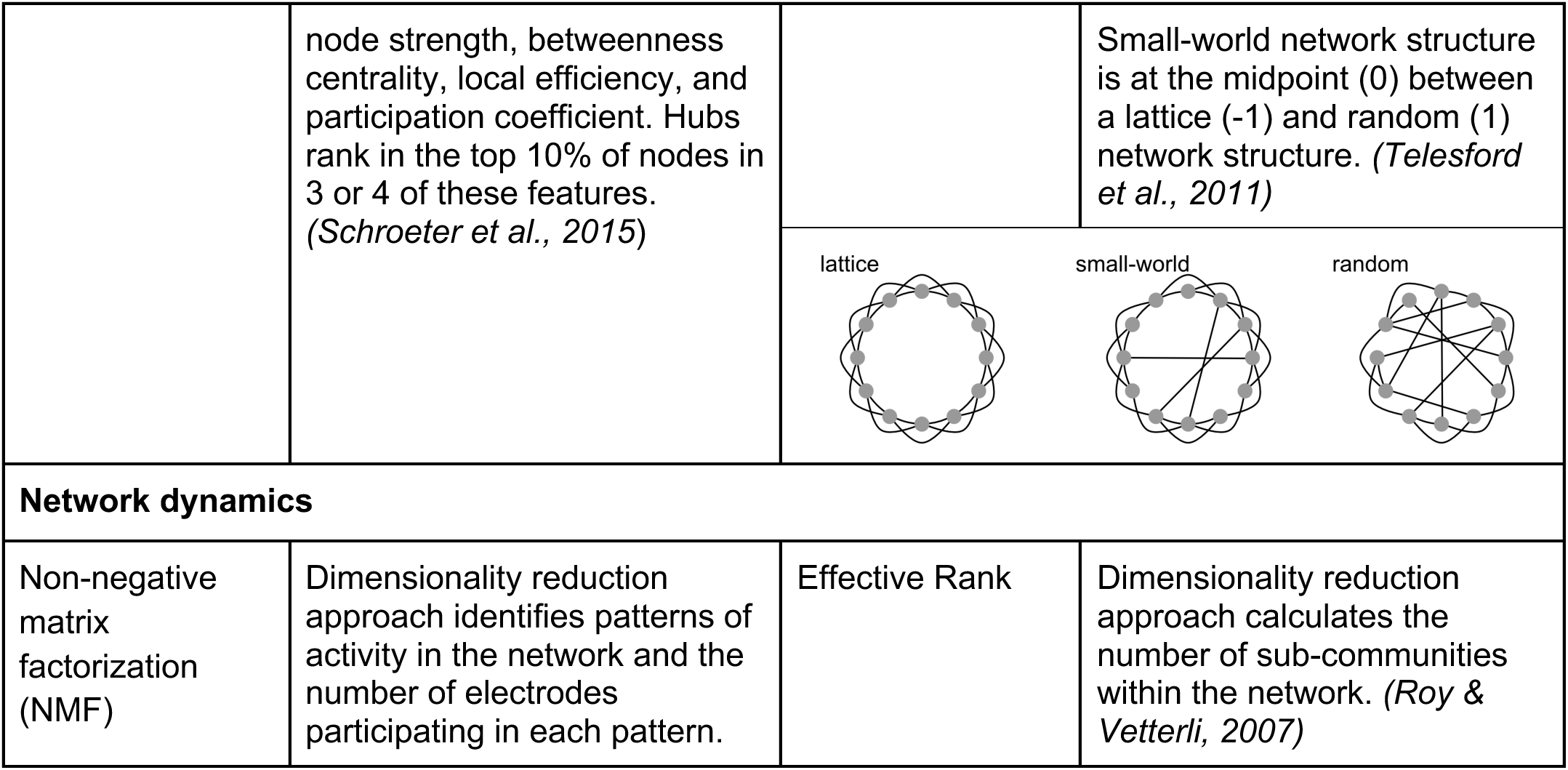
Functional connectivity, network topology, and network dynamic metrics in MEA-NAP.

**Figure 3.**
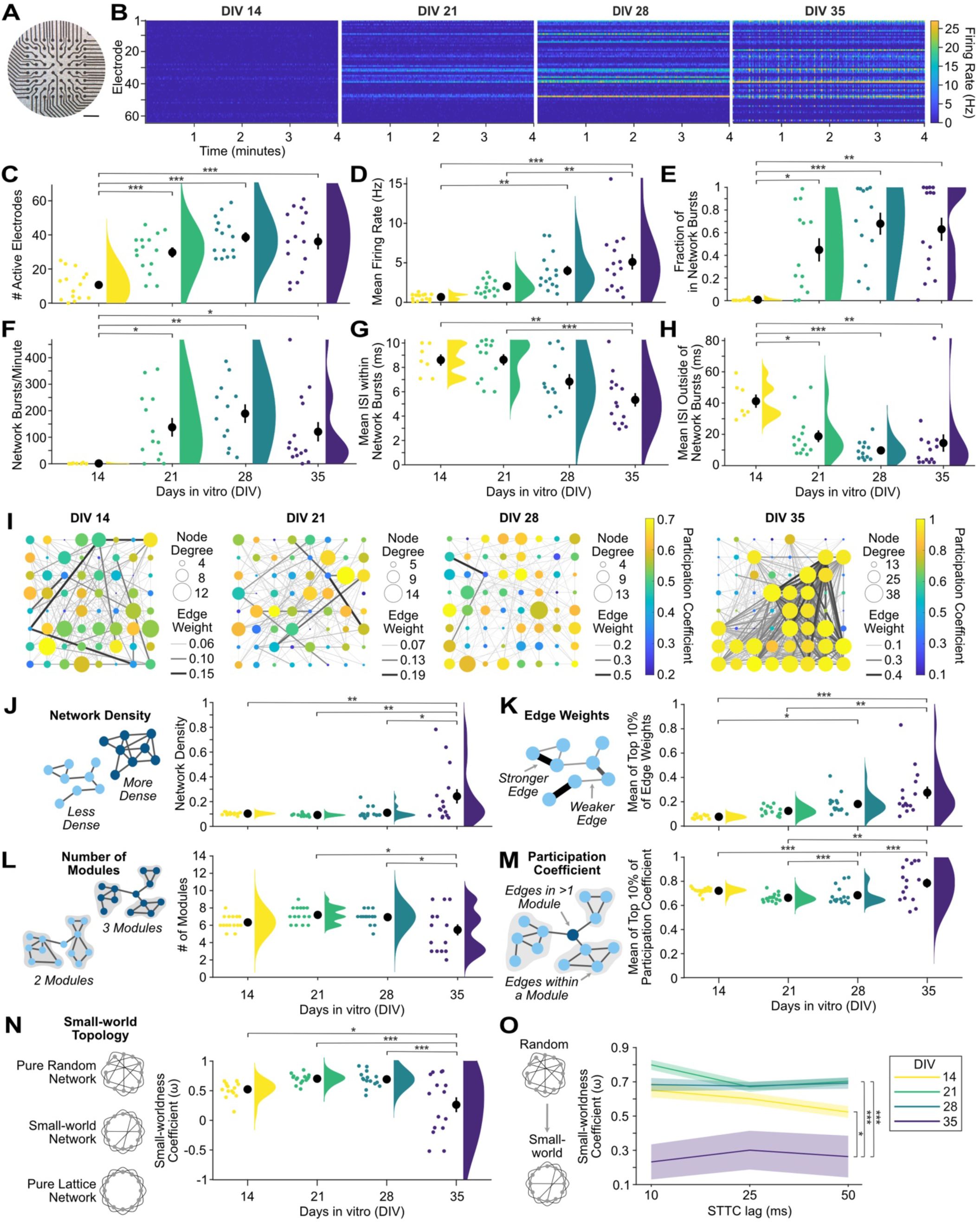
Human 2D NGN2 iPSC-derived cortical cultures develop microscale functional networks. **A.** Photo of NGN2 neurons on an Axion 64-electrode MEA. Scale bar 0.5 mm. **B.** Raster plots of the last 4 minutes of representative 10-minute MEA recordings from days-in-vitro (DIV) 14, 21, 28 and 35 from the same culture show increase in firing and burst rates. **C-H.** Scatter plots with mean ± SEM (black circles & lines) and density curves show a developmental increase in number of active electrodes (**C**), mean firing rate (**D**), fraction of in-network bursts (**E**), and network burst rate (**F**) and a decrease in the mean ISI within (**G**) and outside (**H**) of network bursts (n=15 cultures). **I.** Representative graphs of the development of microscale networks from DIV 14-35 from a single culture. Node strength (circle size), edge weight (line thickness) and density (number of lines), and participation coefficient (node color) increased over development. **J-L.** Illustrations (left) of each network metric with nodes (blue circles) and edges (lines). Scatter plots (right) for each metric with mean ± SEM and density curves show a developmental increase in the network density (**J**) and mean of the top 10% of edge weights (**K**). The number of modules (**L**) decreased, while the mean participation coefficient of the top 10% of nodes (**M**) increased. **N.** Scatter plots with mean ± SEM and density curves show the small-worldness coefficient (ω). Values near 1 represent more randomly connected networks, while values approaching 0 indicate small-world networks and −1 lattice-like (highly connected) networks. **O.** Line graph of mean (lines) ± SEM (shading) for the small-worldness coefficient (ω) at DIV 14-35 as calculated with three different spike time tiling coefficient (STTC) lags (10, 25, 50 ms). Statistical significance was determined with a one-way ANOVA and the Tukey-Kramer method to adjust for multiple post-hoc comparisons. * p<0.05, **p<0.01, *** p<0.001

### Node cartography identifies the development of hub nodes in microscale functional networks

To examine the roles of neurons, or groups of neurons, in the network activity, we applied node cartography ^20^ (Figure 4A) in MEA-NAP. This method uses the within-module z-score and the participation coefficient for each node to distinguish between hubs, non-hubs and peripheral nodes (Figure 4B). Hub nodes connect local clusters, modules, or subcommunities and facilitate participation in the overall network activity ^21^. Hub nodes exist at the brain-region ^5^ to microscale ^7^ levels and have been shown, for example in in vivo microscale murine hippocampal networks, to be stable over time using two-photon calcium imaging ^22^. Importantly, functional magnetic resonance imaging (fMRI) of brain networks reveals that hub nodes are critical for efficient task performance at the cognitive level, allowing for information processing that is integrative across the whole network and segregated into subcommunities ^23^. To determine hub and non-hub roles in neuronal cultures, the role designation boundaries are automatically determined in MEA-NAP using a density landscape based on the distribution of the within-module z-score and the participation coefficient values for the entire dataset (Figure 4C). We applied node cartography to MEA recordings from 2D murine cortical cultures (n=19) revealing both hub and non-hub nodes (Figure 4D) and a developmental increase in kinless hub and non-hub neurons from DIV 14-28 (Figure 4E and 4F, p<0.001, one-way ANOVA, with post-hoc paired t-tests for age). This example in wild-type mice illustrates how node cartography in MEA-NAP could be used in murine disease models to identify developmental differences in hub nodes. This may be particularly important for elucidating microscale functional network defects in genetic causes of autism spectrum disorder and intellectual disability.

**Figure 4.**
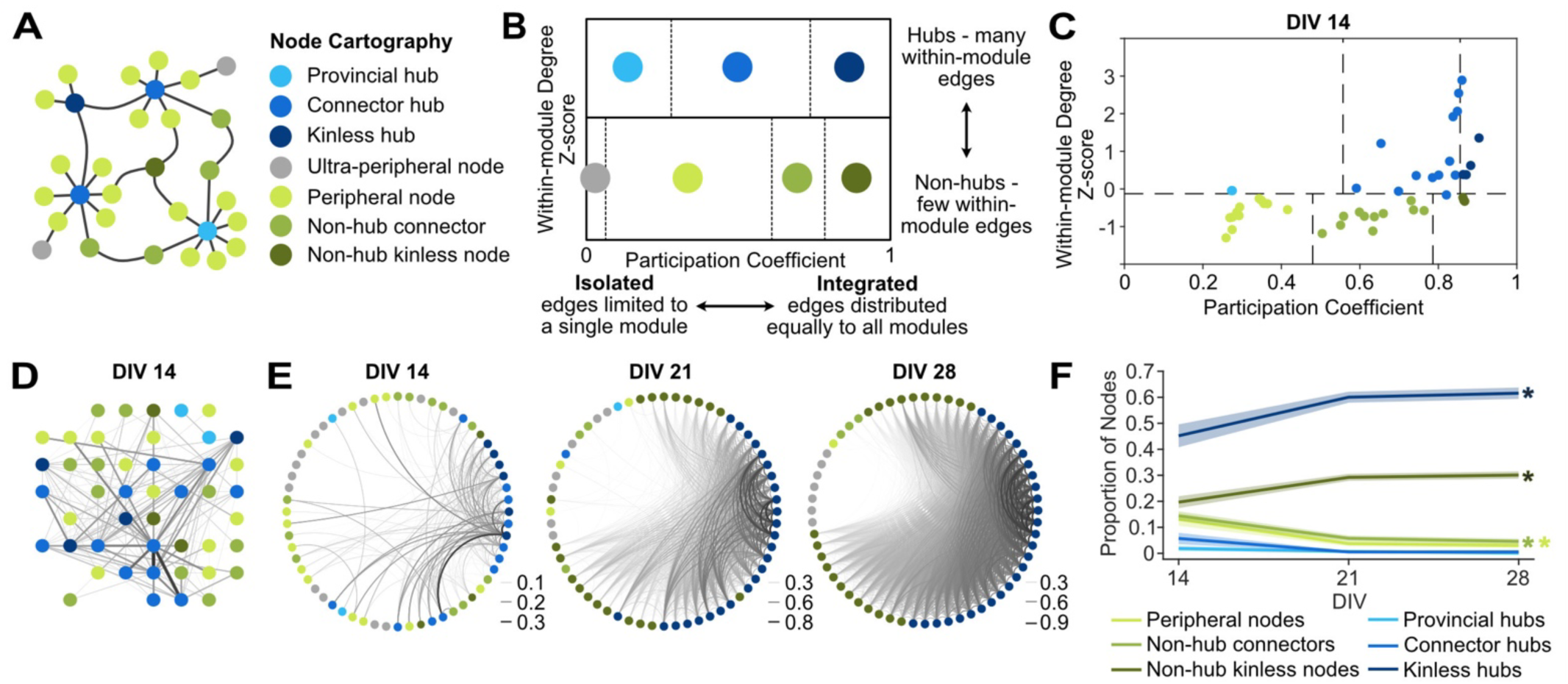
Node cartography reveals the roles of individual nodes in network activity. **A.** Schema of a network shows 7 different node cartography roles including hub (blue) and non-hub (green and gray) designations. **B.** Diagram shows how node cartography roles (colors) are determined based on the within-in module degree z-score and participation coefficient. **C.** Scatter plot of node cartography (color) for a representative 2D mouse cortical culture at days-in-vitro (DIV) 14. Boundaries (dashed lines) for determining hub versus non-hub and subtypes were automatically set for the experimental dataset in MEA-NAP using the density landscape (not pictured). **D.** Network plot with node cartography (color) in the spatial arrangement of the MEA. Nodes (circles) and edges (lines) show functional connectivity for the representative culture in C. **E.** Circular network plots show development of node cartography (colors) and functional connectivity (lines show edges) for representative 2D mouse cortical culture from DIV 14-28. Line thickness represents edge weights (scaled for each DIV) to show modularity. **F.** Comparison of the proportion of node cartography roles (colors) for the entire dataset of 2D mouse cortical cultures (n=19) from DIV 14-28. Mean (solid lines) ± SEM (shading). There were significant effects of age (* p<0.001, one-way ANOVA).

### Identification of subcommunities within microscale functional networks based on the network dynamics

Dimensionality reduction methods can detect patterns of activity observed in multiple nodes within the network ^24^ and can reveal population-level effects of age or condition not apparent from pairwise metrics used elsewhere in MEA-NAP. Dimensionality reduction approaches are commonly applied to multi-unit recordings of neuronal activity in in vivo animal models and can capture complex population-level responses on cognitive tasks ^25^. Thus, this approach can also provide a link between network development and disease perturbations observed in vitro through MEA recordings of neuronal cultures and relevance of these features for microscale information processing at the macroscale level in vivo. Dimensionality reduction also provides a complementary approach to the metrics based on pairwise comparisons in MEA-NAP, such as number of modules, within-module z-score, and participation coefficient for identifying subcommunities in the microscale networks.

To identify the participation of neurons near individual electrodes in functional subnetworks in MEA recordings, MEA-NAP applies non-negative matrix factorization (NMF). This approach has been applied at the macroscale to fMRI in humans, where the participation of individual nodes in multiple subnetworks correlated with task-based flexibility and performance ^26,27^, and at the microscale to in vivo MEA recordings in rat, where NMF detected the onset and spread of induced epileptic activity ^28^. We utilized NMF to compare the development of subcommunities in 2D murine cortical (n=19) and hippocampal (n=10) cultures (Figure 5). To illustrate how NMF reveals differences in the patterns of activity in MEA recordings, we first selected representative raster plots from MEA recordings of DIV 21 cortical and hippocampal cultures with similar firing rates (Figure 5A). Although both raster plots show network bursts, the raster plots of the top three NMF components reveal different patterns of activity underlying the bursting activity (Figure 5B). The NMF components are ranked based on the percent of the variance observed in the MEA recording. MEA-NAP quantified the number of NMF components in two ways. First, MEA-NAP calculated the number of NMF components that explained 95% of the variance observed in the MEA recordings was quantified (Figure 5C). Second, the number of significant NMF components was determined by comparing the number of NMF components from a random spike time matrix created by shuffling the time series in the original recording (Figure 5D). These complementary methods allow comparisons of the number of dominant (or major) NMF components identified in the MEA recordings and the total number of components. This may be particularly important, for example, in disease models where bursting activity is preserved or enhanced leading to a similar number of dominant patterns but activity of subgroups or modules in the network (activity observed outside of the bursts) is lost. The 2D murine cortical cultures showed a trend toward a higher mean number of significant NMF components than hippocampal cultures (Figure 5E) despite similar (DIV 14)---or even smaller (DIV 21-28)---mean network size (Figure 5F and 5G). Thus, NMF analysis revealed that cortical cultures may support a larger number of patterns of activity than hippocampal cultures, in which the firing of action potentials outside of network bursts becomes less frequent over early development (DIV 14-28) compared to cortical cultures ^29^. NMF analysis can facilitate age and/or experimental condition comparisons of the ability of microscale functional networks to support a diversity of patterns.

**Figure 5.**
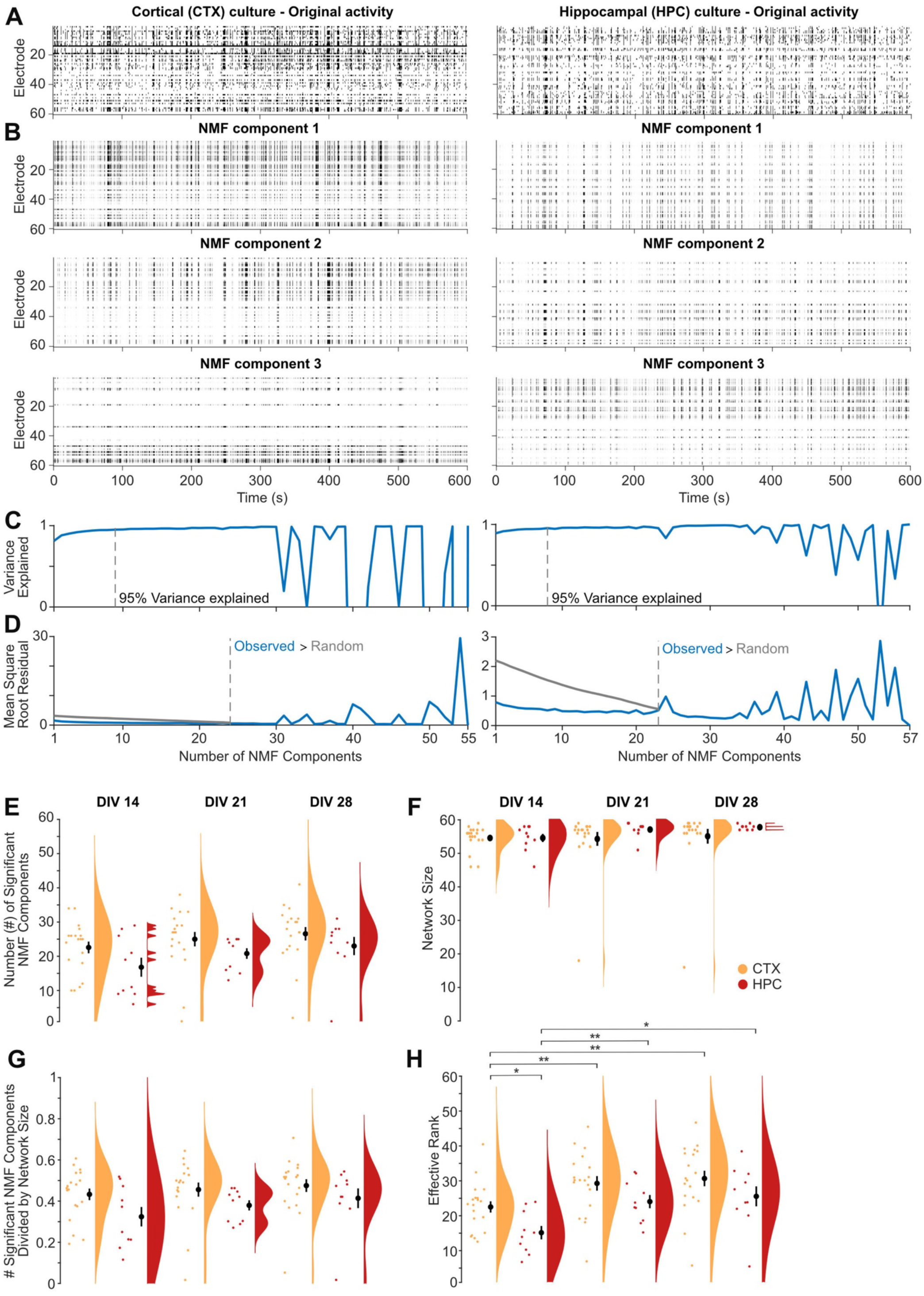
Dimensionality reduction approaches reveal fewer subcommunities in 2D mouse hippocampal than cortical cultures. **A.** Raster plots of representative MEA recordings from 2D primary cortical (left) and hippocampal (right) murine cultures at DIV 21 show action potential firing rate (binned by 100 ms) by electrode (rows) for 10 minutes of spontaneous activity. **B.** Raster plots of the top three non-negative matrix factorization (NMF) components for the MEA recordings in A. **C.** Number of NMF components that explained 95% of variance (dashed gray line) in the MEA recordings in A. **D.** Comparison of the mean square root residual by the number of NMF components for the MEA recordings in A (blue line) and the MEA recordings shuffled. The number of significant components (dashed line) was determined where the mean square root residual from the observed recordings was greater than that from the shuffled recordings (random). **E-H.** Scatterplot, mean ± SEM, and density curves for number of NMF components (E), network size (F), number of NMF components normalized by network size (G), and effective rank (H) comparisons by age and cortical (CTX, orange, n=19) versus hippocampal (HPC, red, n=10) cultures.

MEA-NAP also applies effective rank ^30^ to MEA recordings of neuronal activity. Effective rank provides a continuous measure of the dimensionality of the population activity. Thus, it bypasses the need for heuristics, such as the elbow method, that are commonly used to determine the number of significant dimensions in principal components analysis (PCA). In contrast to the NMF analysis, effective rank provides a single number representing the number of patterns or subcommunities within the network. An effective rank of one indicates fully correlated activity across all nodes in the network, while an effective rank equal to the number of recorded nodes indicates independent activity in all nodes. Effective rank can be compared across networks of different size (e.g., over development in vitro) by dividing it by the number of active electrodes, what we have termed relative effective rank. In the murine 2D cultures, the mean effective rank was higher in cortical than hippocampal cultures at DIV 14 and 21 (Figure 5H). This finding was consistent with the NMF analysis that identified more patterns of activity in the cortical cultures. Computational modeling suggests that lower effective rank in networks may improve learning dynamics compared to higher effective rank ^31^.

## Discussion

### Main findings

Here we provide evidence that MEA-NAP can be exploited as a user-friendly computational tool for batch analysis of MEA data revealing functional connectivity, network topology and network dynamic features in 3D brain organoids and 2D human-derived or murine neuronal cultures. MEA-NAP includes a user-friendly GUI and detailed documentation to assist users new to MATLAB, MEA analysis, and/or network science. MEA incorporates leading methods for robust spike detection, including template-based methods, and for determining functional connectivity, including combining the spike time tiling coefficient with probabilistic thresholding to identify significant connections. MEA-NAP incorporates many graph theoretical metrics from the Brain Connectivity Toolbox and other sources along with state-of-the-art methods for normalizing network metrics. MEA-NAP integrates null models to determine significant network features including the development of small-world topology. MEA-NAP reveals the functional connectivity and network topology in 3D human cerebral organoids and 2D human NGN iPSC-derived cortical cultures. MEA-NAP introduces node cartography for MEA recording and adapts the density landscape approach to automatically determine hub role boundaries for each dataset. This approach reveals an increase in hub nodes and integration of nodes into the overall network activity in murine cortical cultures over early development Dimensionality reduction methods in MEA-NAP including NMF and effective rank reveal subcommunities of neurons within the networks by their patterns of activity observed in multiple electrodes that were greater in cortical than hippocampal murine cultures.

### Comparison with other publicly available tools

MEA-NAP incorporates methods for analyzing functional connectivity, network topology, and dimensionality reduction from multiple disciplines of network science and adapts these methods for robust application to microscale functional networks in MEA recordings (Tables 1 and S1). We compared MEA-NAP’s performance on the dataset from Schroeter et al. (2015). MEA-NAP recapitulates the development graph theoretical metrics in these 2D hippocampal cultures (Figure S1). MEA-NAP expands the offering beyond current publicly available MEA and network toolboxes (Tables S2 and S3) through creating a streamlined approach for batch analysis from comparison of spike detection methods (including template-based methods) to inferring significant functional connections and comparing network topological and dynamic features. MEA-NAP has been adapted for and tested on multiple MEA systems (Multi-channel Systems and Axion Biosystems) and on recordings from 2D and 3D murine or human-derived neuronal cultures. MEA-NAP automatically generates the figures needed to examine both the validity and the age and genotype comparisons. This is particularly helpful for users new to MATLAB and/or network science. MEA-NAP also facilitates comparisons with other microscale network approaches. In vivo calcium imaging of the rodent hippocampus has identified hubs that are stable over time. This approach provides single-cell resolution and can connect network topological features to task performance ^22,32^. Functional connectivity and network topology analysis of MEA recordings provide a complementary approach. MEA recordings have a higher temporal resolution necessary for detecting action potentials, and late embryonic and early postnatal time points in network development can be studied before in vivo imaging is practicable.

### Limitations

MEA-NAP is designed for 60-electrode (Multi-channel Systems) or 64-electrode (Axion Biosystems) microelectrode arrays. Although advanced MATLAB users can adapt the code to accommodate other electrode numbers, spatial arrangements, and/or spike-sorted data from in vivo recordings (e.g., Neuropixel), multiple topographical features may not work on MEAs with too few electrodes (e.g., 12 or 16) and the computational time necessary to determine functional connectivity in higher numbers of electrodes (e.g., 384 electrodes or high-density arrays) may be too long. The default parameters set for the spike detection (CWT, cost parameter), burst detections, and functional connectivity (lag) have been tested on a variety of datasets from 2D and 3D, mouse and human, different disease models. However, the user will need to confirm that these parameters are performing well on their data. We have tested MEA data acquired at 12.5 and 25 kHz for 5 to 12 minutes. Longer recording times may require more computational time and/or resources.

Spikes detected at individual electrodes may originate from one (single unit) or more (multi-unit) neurons near each electrode. MEA-NAP uses this multi-unit activity. The network features compared in MEA-NAP are seen across spatial scales in the brain. We chose not to incorporate spike sorting, as it is difficult to establish a ground truth to validate even the leading spike sorting methods on one’s own experimental data. For determining significant functional connections, the spike time tiling coefficient (STTC) is in most cases rate-independent ^11^; however, the choice of STTC lag can alter downstream graph metrics. Thus, the lag comparison tools in MEA-NAP should be used. Probabilistic thresholding is less likely to introduce false positives or negatives than absolute or proportion thresholding ^14^. We have incorporated leading methods for determining significance in multiple network metrics; however, adjustment of the p value for multiple comparisons should be considered based on the experimental design ^33^. Users are encouraged to perform pre-power analysis and “mask out” any comparisons with insufficient power ^34^.

### Broader Application

We created MEA-NAP to facilitate application of network-level analysis to MEA recordings performed by neurobiologists studying in vitro models of neuronal development and disease. In addition, our data provides an extensive reference resource for neuronal network behavior studies. MEA-NAP can reveal how perturbations such as genetic mutations in human iPSC-derived or mouse models impact functional connectivity, network topology and network dynamics at the microscale. Thus, MEA-NAP can also inform mechanistic and therapeutic studies in vitro. Future application of MEA-NAP to microscale functional networks observed in MEA recordings from patient iPSC-derived neurons can be compared to application of graph theoretical metrics in functional imaging or EEG students in the same patient or patient populations. Combining these approaches could decipher the missing link between microscale and macroscale brain development and dysfunction in neurologic and psychiatric disorders ^35^.

## Methods

### Murine 2D cultures

#### Cell culture initiation

The MEA recordings and methods for the murine primary hippocampal cultures were previously published ^8^. For the murine primary cortical cultures, neonatal C57BL/6J mice were sacrificed at postnatal day 0 in accordance with UK Home Office regulations by first inducing hypothermia prior to decapitation. The heads were quickly submerged in sterile ice-cold phosphate buffer solution (PBS, Gibco, 14190094), and the cortices were dissected in sterile ice-cold PBS under a stereoscope followed by removal of the meninges. The cortices were next chemically dissociated using a 1:1 mixture of papain (Sigma, P5306) and sterile PBS at 37 °C for 25 minutes. The reaction was stopped by adding neurobasal media (Gibco, 21103-049) with B27 supplement (NB-B27; B27 supplement, Gibco, 17504-044) with 4% fetal bovine serum (Gibco, 175004044) and the cortices were manually dissociated using a pipette. A small volume (20 μL) of the cells were removed for manual cell counting with a hemocytometer and the remaining volume was then centrifuged for 10 minutes at 0.4 r.c.f. The supernatant was removed, and the pellet was resuspended in neurobasal media with the B27 supplement (NB-B27). The cells were resuspended in the volume necessary to ensure that 15 μL would contain 5×10^4^ cells, which were plated on single-well 60-channel MEA chips (Multi-channel systems, 60MEA200/30iR-ITO-gr).

#### MEA preparation and plating

The MEA chips were treated in advance with heavy-weight poly-L-lysine (PLL; Sigma, P4832) for 5 minutes up to 24 hours at 37 °C in the incubator, followed by three PBS washes to ensure full removal of the PLL, before adding 7 μL of laminin (Sigma, L2020) directly over the MEA grid. A 15 μL aliquot of 5×10^4^ cells was added directly to the laminin on the MEA grid, and the MEA chips were incubated for at least 20, but not more than 30, minutes. Visual inspection under a light microscope was used to confirm cell adhesion prior to adding 300 μL of NB-B27 at 37 °C to the MEA well. The MEA cultures were incubated overnight. The following day an additional 300 μL of NB-B27 with 0.25% Glutamax (Gibco, 35050-038) was added to each well. Cultures were maintained in the incubator at 37 °C with humidity control and 5% carbon dioxide. The 30% media (180-200 μL/well) was exchanged three days per week with fresh NB-B27 with 0.25% Glutamax at 37 °C. MEA recordings were made weekly from DIV 7-35 using the MEA2100 system (Multi-channel systems).

### 2D human iPSC-derived cortical cultures

#### Cell culture initiation

Frozen DIV 4 stock of NGN2 iPSC-derived cortical neurons (Brigham & Women’s Hospital Neuro iHub) were gently thawed and resuspended in NB-B27 media with RHO/ROCK pathway inhibitor (RI, 10 μM, Stemcell Technologies, Y-27632). A small volume (10 μL) of the cells was removed for automated cell counting (Invitrogen, Countess 3). The remaining volume was centrifuged for 5 minutes at 0.4 r.c.f. The supernatant was removed, and the pellet was resuspended in the volume of NB-B27 with supplements (NB-B27-S) and RI necessary to ensure that 15 μL would contain 35×10^3^ cells, which were plated on sterile 6-well (64-channels per well) CytoView MEAplates (Axion Biosystems, M384-tMEA-6W). The NB-B27-S included human ciliary neurotrophic factor (CNTF, 10 ng/mL, 1:1000, Peprotech, AF-450-13), glial-derived neurotrophic factor (GDNF, 10 ng/mL, 1:1000, Peprotech, 450-10), brain-derived neurotrophic factor (BDNF, 1:1000, 10 ng/mL, Peprotech, 450-02) recombinant proteins, Glutamax (1%, Gibco, 35050), MEM Non-Essential Amino Acids Solution 100X (0.5%, Invitrogen, 11140-050), doxycycline hyclate (DOX, 2 μg/mL, Sigma, D9891), and dextrose 20% (1.5%, Sigma, D9434).

#### Astrocyte preparation

Frozen normal human astrocytes (CC-2565 NHA-Astrocytes AGM, Cryo amp, Lonza, 185115) were gently thawed in AGM Astrocyte Growth Medium BulletKit (Lonza, CC-3186) and automated cell counting performed. The astrocytes were centrifuged for 5 minutes at 0.4 r.c.f. The supernatant was removed, and the pellet was resuspended in the volume of NB-B27-S with RI necessary to ensure that 20 μL would contain 12×10^3^ cells.

#### MEA preparation and plating

The MEAs in the 6-well plates were treated in advance with polyethylenimine (PEI) for 12-24 hours at 4 °C (or heavy-weight PLL for 5 minutes up to 24 hours at 37 °C in the incubator), followed by three PBS washes to ensure full removal of the PEI or PLL, before adding 12 μL of laminin directly over each MEA grid. The 15 μL aliquot of 35×10^3^ cells was added directly to the laminin on the MEA grid. The MEA chips were incubated for at least 20, but not more than 30, minutes. During the incubation, the astrocytes were prepared. The MEA plates were removed briefly one at a time to plate 20 μL of the astrocyte solution (12×10^3^ cells) per well, distributed in 3 or 4 5-7 μL drops around– but not touching–the drop of neurons on the grid. After the incubation of the neurons was complete, cell adhesion was confirmed with visual inspection under a light microscope prior to adding 400 μL of NB-B27 with the supplements and RI at 37 °C to each MEA well. The MEA cultures were incubated overnight. The following day an additional 350 μL of NB-B27 with the supplements—without RI—was added to each well. Cultures were maintained in the incubator at 37 °C with humidity control and 5% carbon dioxide. Three days per week ∼30% of media was exchanged with fresh NB-B27 with CNTF, GDNF, BDNF, and DOX at 37 °C. The volume of media per well was gradually increased to 1.5 mL per well by DIV 21. MEA recordings were made weekly using the Maestro MEA system (Axion Biosystems).

### 3D human air-liquid interface cortical organoids

Human air-liquid interface cerebral organoids (ALI-COs) were generated from the H9 embryonic stem cell line using published methods ^3,9^. In brief organoids were sliced between DIV 55-60 and cultured at the air-liquid interface. On the day of recording, the ALI-COs were transferred from the membrane to the 60-channel 3D MEA chip (Multi-channel systems, 60-3DMEA250/12/100iR-Ti-gr) filled with neurobasal media with B27 supplement. The ALI-COs were positioned on the grid, and a platinum harp with nylon strings was used to gently hold the ALI-CO in place. A lid with a semi-permeable membrane (ALA Science, MEA-MEM5) was placed on the MEA chip to reduce evaporation. The chip was transferred to the MEA2100 system. After MEA recording, the ALI-COs were either fixed for immunohistochemistry or discarded.

### MEA data acquisition

#### Recordings using MCS MEA systems

Data for the 2D mouse hippocampal cultures was acquired with an MEA1600 MEA System (Multi-channel systems) as previously described ^8^. The 2D mouse cortical culture and the 3D human cerebral organoid cultures were recorded with an MEA2100 dual headstage MEA system (Multi-channel systems), with the temperature controller (Multi-channel systems, TCX-2) set to 37 °C. Raw data was acquired at 25 kHz for 5-12 minutes using the MCRack software (Multi-channel systems). Raw data was exported using MCTool (Multi-channel systems) and converted to MATLAB format (.mat) using custom scripts included in MEA-NAP. All photographs of the cultures on the MEA grid were taken with a standard digital camera attached to a light microscope using a scope attachment and saved in .jpeg format.

#### Recordings using Axion Biosystem MEA system

Data for the 2D human iPSC-derived cultures was recorded with a Maestro MEA system, with the temperature set to 37 °C. Raw data was acquired at 12.5 kHz for 10 minutes using the Axis software (Axion Biosystems). Raw data was exported to MATLAB format (.mat) using custom scripts, including publicly available MATLAB scripts from Axion for separating the 6 MEA recordings per multi-well plate into individual MEA recordings. All photographs of the cultures on the MEA grid were taken with an iPhone (Apple) through the scope of an inverted light microscope and were exported in .jpeg format.

### MEA-NAP data analysis

All MEA data was analyzed using our MEA network analysis pipeline (MEA-NAP) in MATLAB (2021b) on a computer with at least 16GB RAM. The pipeline inputs included: (1) raw voltage traces acquired with either a 60-channel MEA1600 or 2100 system (Multichannel Systems) or 64-channel each multi-well MEA plates (Axion Biosystems) and (2) a batch analysis file (.csv) with the filenames to be analyzed and the age and group information. Step-by-step methods for MEA-NAP are available in our online documentation at https://github.com/SAND-Lab/MEA-NAP/ (and permanently at https://doi.org/10.7910/DVN/Z14LWA).

#### Data conversion tools and filtering

Voltage time series collected with the MEA2100 or MEA1600 system (Multi-channel systems) were converted to MATLAB format using the MC_Tool (Multi-channel systems). Voltage time series collected with the Maestro (Axion Biosystems) 6-well plates were converted to MATLAB format using a custom script included with MEA-NAP. We provide a sample MEA dataset in MATLAB format (Supplemental Resource 1). We also include MATLAB functions to facilitate the import of raw MEA data from MCS or Axion Biosystems MEA systems. In MEA-NAP, the voltage time series are first bandpass filtered (third-order Butterworth filter, 600-8000 Hz).

#### Spike detection and validation tools

Spike detection was performed using multiple methods. (1) Template-based spike detection was performed using the continuous wavelet transform (CWT) ^12,13^ with MATLAB BiorSplines family bioorthogonal wavelets (bior1.5, bior1.3) and Daubechies orthogonal wavelet (db2) and a cost parameter of −0.12. (2) Threshold-based spike detection using the mean absolute deviation (MAD) of the voltage signal ^36^ with thresholds of 3, 4, and or 5 MAD. Plots generated for each recording included a running average of the spike frequency for the entire MEA for each spike detection method, an overlay of 50 sampled spikes and the averaged waveform(s) of spikes detected for each method for a subset of electrodes, and sample voltage traces with spikes detected by method from 9 electrodes. These plots were reviewed for each MEA recording to compare the performance and confirm accuracy spike detection.

#### Spike features and network burst metrics

The spike frequency for each electrode was calculated as the number of action potentials per second (raster plot) divided by the length of the recording (MEA heatmap, scatter plot with mean and density curve) for each recording. The number of active channels was calculated as the number of electrodes per MEA in which the number of spikes detected divided by the length of the recording was greater than 0.01 Hz (or 0.1 Hz for the NGN2 dataset, Figure 3). The mean and median spike frequency were calculated for each recording and compared by age and group. Here, network bursts were defined as a minimum of 10 spikes in at least 3 channels with the ISI_N_ threshold ^18^ set automatically; these parameters are easily adjustable in MEA-NAP in the advanced settings. Network bursts were characterized by rate (number of network bursts divided by length of the recording in minutes), mean number of electrodes (channels) participating in the network burst, mean network burst length (seconds), mean inter-spike interval (ISI) within and outside (between) network bursts (both in milliseconds), coefficient of variation in the inter-network burst intervals, and the fraction of single-channel bursts that occur in network bursts for each recording. These features were automatically plotted as scatter plots with mean, SEM, and density curves for age and group comparisons.

#### Inferring significant functional connections

Functional connectivity was inferred through pairwise comparisons of the spike trains with the spike time tiling coefficient (STTC) ^11^. Time lags for the STTC were compared for 10, 25 and 50 milliseconds for most recordings (see Supplemental Figure S1H). Significant functional connections were determined using probabilistic thresholding ^14^. For each pair of electrodes, a circular shift was performed on one of the spike trains, and the STTC was calculated for 180 iterations. Significant functional connections were identified where the real STTC value exceeded the 95th percentile of the STTC values for the 180 different circular shifts. These parameters are easily adjustable in MEA-NAP. The effect of the number of iterations on the threshold for determining significant edges was reviewed. Weighted adjacency matrices with only the significant functional connections were used for subsequent analysis of functional connectivity and network topology metrics, with the exception of the dimensionality reduction approaches which did not rely on pairwise comparisons. For the network metrics and statistical comparisons in Figures 2 and 3, a STTC lag of 50 ms was used for the human iPSC-derived organoid and monolayer cultures. For the network metrics and statistical comparisons in Figures 4 and 5, a STTC lag of 10 ms was used for the postnatal mouse cortical and hippocampal postnatal cultures.

#### Graph theoretical metrics

The graph theoretical metrics calculated in MEA-NAP are illustrated in Table 1 and the source code referenced in Table S1. The node degree, edge weight, node strength, network size, network density, and global efficiency were calculated using dedicated functions from the Brain Connectivity Toolbox (BCT) ^6^. To determine the local efficiency, the weight_converstion function from the BCT was used first for normalization followed by the efficiency function from the BCT. Modularity (score and number of modules) were derived from a consensus clustering method ^37^ to produce modularity groups. The within-module z-score was calculated using the module_degree_zscore function from the BCT with the modularity affiliation vector calculated above as an input argument. The betweenness centrality (BC) was calculated using the BCT function. The BC was next normalized to the range 0 to 1 using the formula BC/((N-1)(N-2)), where N is the number of nodes in the network. The normalized participation coefficient was calculated using a method described ^38^. The clustering coefficient was calculated using the BCT function and normalized using a lattice model ^39,40^. The characteristic path length was calculated using the BCT function and normalized using a random null model ^39^. The small-world topology was calculated in two different approaches. The small-world coefficient (σ) was calculated by dividing the clustering coefficient, normalized to a random network model, by the characteristic path length, normalized to a random network model ^41^. The small-world coefficient (ω) was calculated by subtracting the clustering coefficient, normalized to a lattice model, from the characteristic path length, normalized to a random network model ^19^. Here, the output was restricted to a range of −1 (pure lattice) to 1 (pure random network) and values near zero represent a small-world network.

#### Node cartography

To define the boundaries for hub and non-hub roles in the node cartography, the density landscape was first calculated using the participation coefficient and within-module z-score for all nodes in the dataset using our custom code based on method ^20^. These boundaries for nodal roles were then applied to individual recordings to classify nodes as ultra-peripheral nodes, peripheral nodes, non-hub connectors, non-hub kinless nodes, provincial hubs, connector hubs, or kinless hubs. The proportion of nodes in each role were compared by STTC lag for individual recording and by age and group between recordings.

#### Dimensionality reduction

Non-negative matrix factorization (NMF) was performed using the built-in MATLAB function. We provided two methods for evaluating the number of components sufficient to describe the observed neural activity. In the first method, we computed the root-mean square residual between the observed activity and the NMF rank-k approximation. We then shuffled the activity of each electrode across time bins to destroy temporal correlations between channels to create a null activity matrix with the same mean activity for each electrode. Then we again computed the residual of the rank-k approximation. The number of components is the largest rank-k approximation such that the residual corresponding to the original activity matrix is lower than for the null activity matrix. In the second method, we found the smallest rank-k approximation such that the variance explained was above 95%. The top 3 components from this rank-k approximation were plotted with the original recording. We also plotted the variance explained of each rank-k approximation and the mean-square-root residual of each rank-k approximation for both the original activity matrix and the null activity matrix. The number of NMF components was compared between recordings and normalized by network size. Effective rank was calculated using methods described in Roy and Vetterli ^30^. Briefly, the effective rank was calculated from the covariance matrix of the downsampled (to 10 Hz) spike matrix. Relative effect rank was defined as effective rank divided by number of active electrodes and could be used to compare between cultures.

#### Data visualization

MEA-NAP automatically generated informative plots for each recording and for age and group comparisons of the entire dataset (see sample output folder in Supplemental Resource 2). The age and group comparison half-violin plots include: a scatterplot of the values for individual nodes or individual recordings (colored circles), the mean ± standard error of the mean (SEM; black circle with error bars), and the density curve (half violin). The density curve represents the probability distribution calculated using kernel density estimation for smoothing. For metrics that have a fixed range (e.g., 0 to 1), the density curve is cut off at the upper and lower limits of the metric. Custom color schemes were created for age and group comparisons using Affinity Designer. MEA-NAP produced the plots in Figures 2-5 in .svg format, which were imported to Affinity Designer. After resizing the individual plots in the combined figures, the size of shapes, line thickness, and scatterplot jitter (in x-axis only) in individual plots were adjusted to a uniform size across all plots, while retaining the x,y center coordinates.

### Statistical analysis

MEA-NAP incorporates statistical tests for evaluating differences between ages and groups. In cases where there is a single group (e.g., wild-type cultures) with multiple repeated observations of the same cultures across ages, we performed repeated measures one-way ANOVA on each recording-level metric (e.g., mean firing rate, mean node strength, effective rank). We also fit the observed recording-level metric with a linear mixed effects (LME) model with age as the main effect and a random effect of the specific culture identity on the intercept. We also performed pairwise t-test comparison by age for each recording-level metric. In cases where only a difference between groups is of interest, we utilized a LME model with the group as the main effect and a random effect of the specific culture identity on the intercept. In cases where cultures of different groups and ages were available, we fit two LME models, one with a main effect of age, group, and a random effect of the culture identity on the intercept, and a second one with an interaction between the two main effects.

Beyond traditional statistical comparisons, we also provide methods for identifying age and genotype differences via classification and regression approaches. By default, we performed classification of either the age or genotype using linear support vector machine, k-nearest neighbor, decision tree, and linear discriminant analysis, with network metrics as features. The models were compared via k-fold cross validation (with the option of stratification), and visualizations were created to compare the model misclassification rate with the baseline chance rate. We also performed regression from network features to obtain age predictions using support vector machine regression, ridge regression, regression tree and feed-forward neural network regression. We compared the mean-squared error between the observed and predicted age using k-fold cross validation. To assess the importance of each feature in the classification process, we performed a process akin to leave-one-out feature selection ^42^. We shuffled the feature of interest across observations and compared the difference in the misclassification rate of the original mode with the shuffled model.

In all cases, the automated statistical analysis step in MEA-NAP is intended to provide an overview of where significant trends in network development or group differences may exist. However, users must check the underlying statistical assumptions, such as normality, for their experimental data are correct. They must also correct for multiple comparisons. For the analysis presented in Figures 3-5 and Supplemental Figure S1, the tabular output files from MEA-NAP were used to perform statistical analysis in MATLAB. For Figures 3 and 4 and Figure S1, a one-way ANOVA was performed to determine whether there was an effect of age (days in vitro) on each metric. Adjusted p-values for multiple pairwise comparisons were calculated using the Tukey-Kramer method. For the network burst rate (Figure 3F), the MATLAB implementation of the Tukey-Kramer method produced unrealistically high p-values; thus, small random numbers were added to the zero values to account for numerical instability. For Figure 5, a two-way ANOVA was performed to determine whether there was an effect of age and/or culture origin (cortex and hippocampal) with the Tukey-Kramer method for post-hoc pairwise comparisons.

## Code availability

All MATLAB code for analysis and production of figures in this paper have been permanently deposited in the Harvard Dataverse: https://doi.org/10.7910/DVN/Z14LWA. The latest version of MEA-NAP can be downloaded at: https://github.com/SAND-Lab/MEA-NAP/

For a full list of previously published code from other sources incorporated in MEA-NAP, please see Table S1.

## Contributions

Concept: R.C.F.,S.B.M.,T.P.H.S.

Development of tools: T.P.H.S., R.C.F., A.W.E.D., J.C., S.B.M., A.B., H.H.S., L.B., E.H., M.H.

Development of documentation: T.P.H.S., E.C., D.O.,S.B.M.

Data generation: G.M.G.,A.D.,M.K.,S.B.M.,Y.Y.

Analysis: T.P.H.S.,D.O.,S.B.M.

Manuscript: S.B.M., T.P.H.S.

Feedback and editing: T.P.H.S.,R.C.F.,J.C.,H.H.S., A.L.,M.A.L.,S.J.E.,O.P.

Funding: S.B.M.,O.P.,A.L.,M.A.L.

Supervision: S.B.M.,O.P., S.J.E., A.L.,M.A.L.,F.D.A.

## Acknowledgments

We would like to thank our funders: the European Commission Horizon 2020 (Marie Skłodowska-Curie Actions Individual European Fellowship, EU Project 700999, S.B.M.), the American Academy of Neurology (Career Development Award, S.B.M.), and the UKRI Medical Research Council (Doctoral Training Partnership, A.W.E.D.). S.B.M. is supported by an NIH NINDS K02 Independent Scientist Award (1K02NS131521-01A1). The A.L. lab is funded by the UKRI Medical Research Council (MRC) Senior Clinical Fellowship Award (MR/X006867/1), which also supported the work of G.M.G. The M.A.L. lab is supported by the UKRI MRC (MC_UP_1201/9). Thank you to the many user testers of MEA-NAP, including undergraduate students in the Synaptic and Network Development (SAND) research group and collaborators from other research groups; animal welfare and husbandry staff at the University of Cambridge; the Neuro iHub at Brigham & Women’s Hospital.

## Supplementary material

**Supplemental Figure S1.**
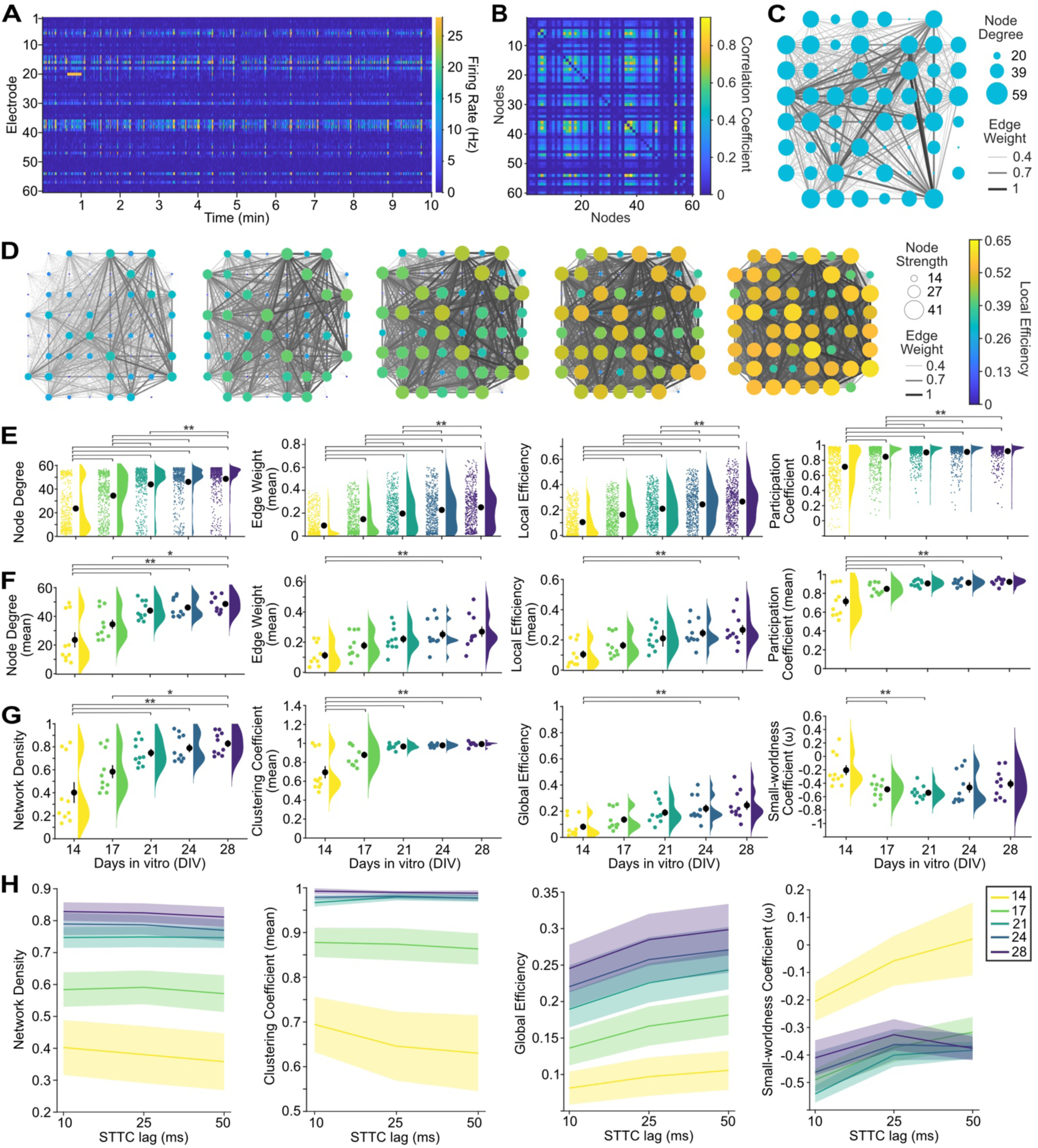
Development of functional connectivity and network topology in 2D murine hippocampal cultures. **A.** Representative raster plot of spontaneous activity in a 10-minute microelectrode array (MEA) recording from primary mouse hippocampal culture. **B.** Adjacency matrix shows correlation coefficient (spike time tiling coefficient, STTC) for significant edges after probabilistic thresholding for recording in A. **C.** Graph of functional connectivity for recording in A. Nodes (circles) represent the activity observed at individual electrodes in the spatial arrangement of the MEA. Number of connections shown as node degree (circle size) and strength of connectivity as edge weight (line thickness). **D.** Development of functional connectivity in representative hippocampal cultures from days-in-vitro (DIV) 14-28 including increase in node strength (circle size), edge weight (line thickness), and local efficiency (circle color). **E.** Comparison of nodal-level network metrics for electrodes (colored circles) from hippocampal cultures (n=10) for node degree, mean edge weight (per node), local efficiency, and participation coefficient. Scatter plots with mean (black circles) ± SEM (error bars) and density curve for DIV 14-28. **F.** Comparison of recording-level network metrics (colored circles) for mean node degree, mean edge weight, mean local efficiency, and mean participation coefficient from DIV 14-28. **G.** Comparison of recording-level network metrics including network density, mean clustering coefficient, global efficiency, and small-worldness from DIV 14-28. **H.** Comparison of recording-level network metrics by STTC lag and developmental age (color, DIV 14-28). Means (lines) ± SEM (shading). For panels E-G, a one-way ANOVA (p<0.01 for all plots) followed by the Tukey-Kramer method to calculate p-values adjusted for multiple post-hoc pairwise comparisons (** p≤0.01, * p<0.05).

**Supplemental Table S1.**
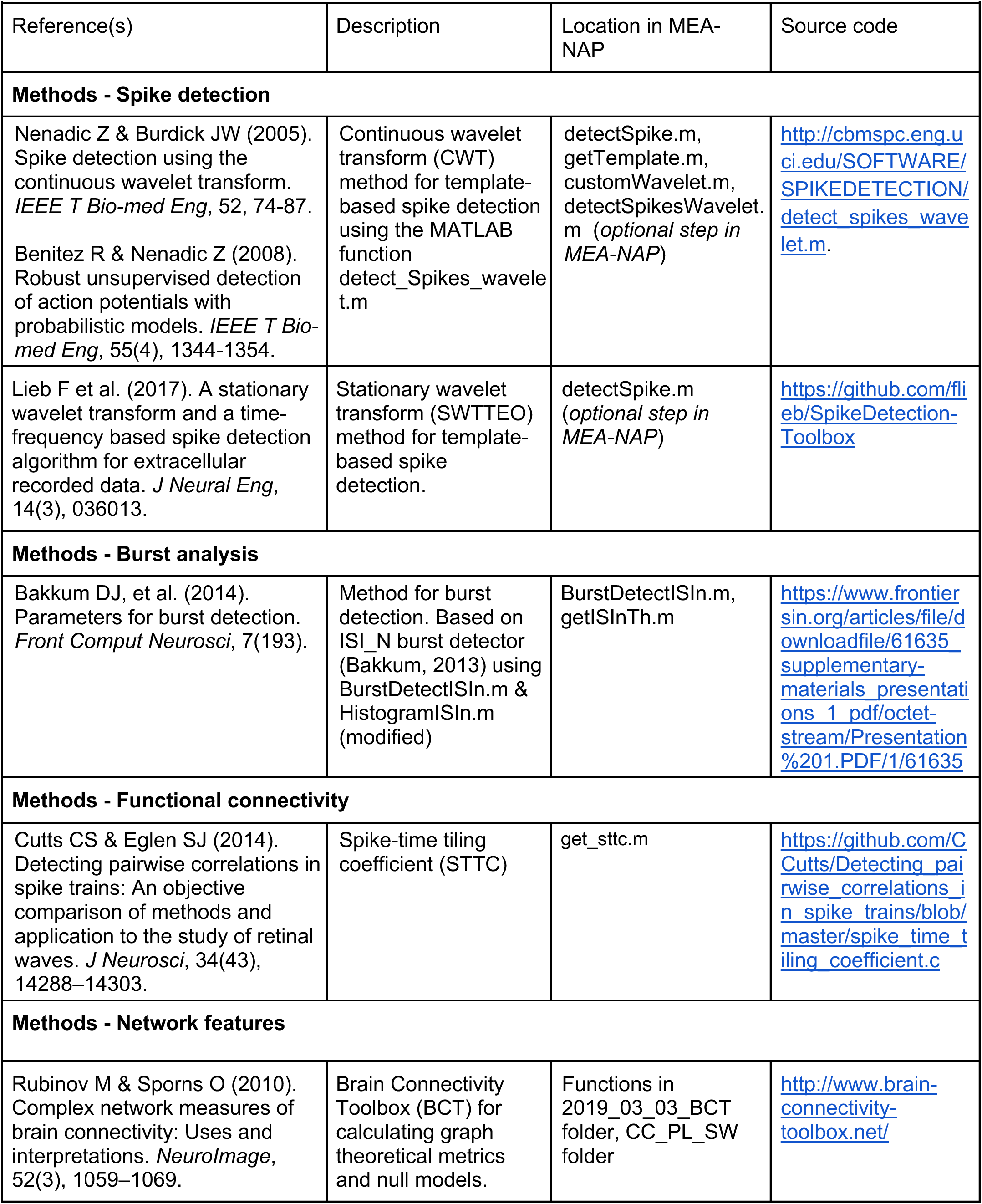
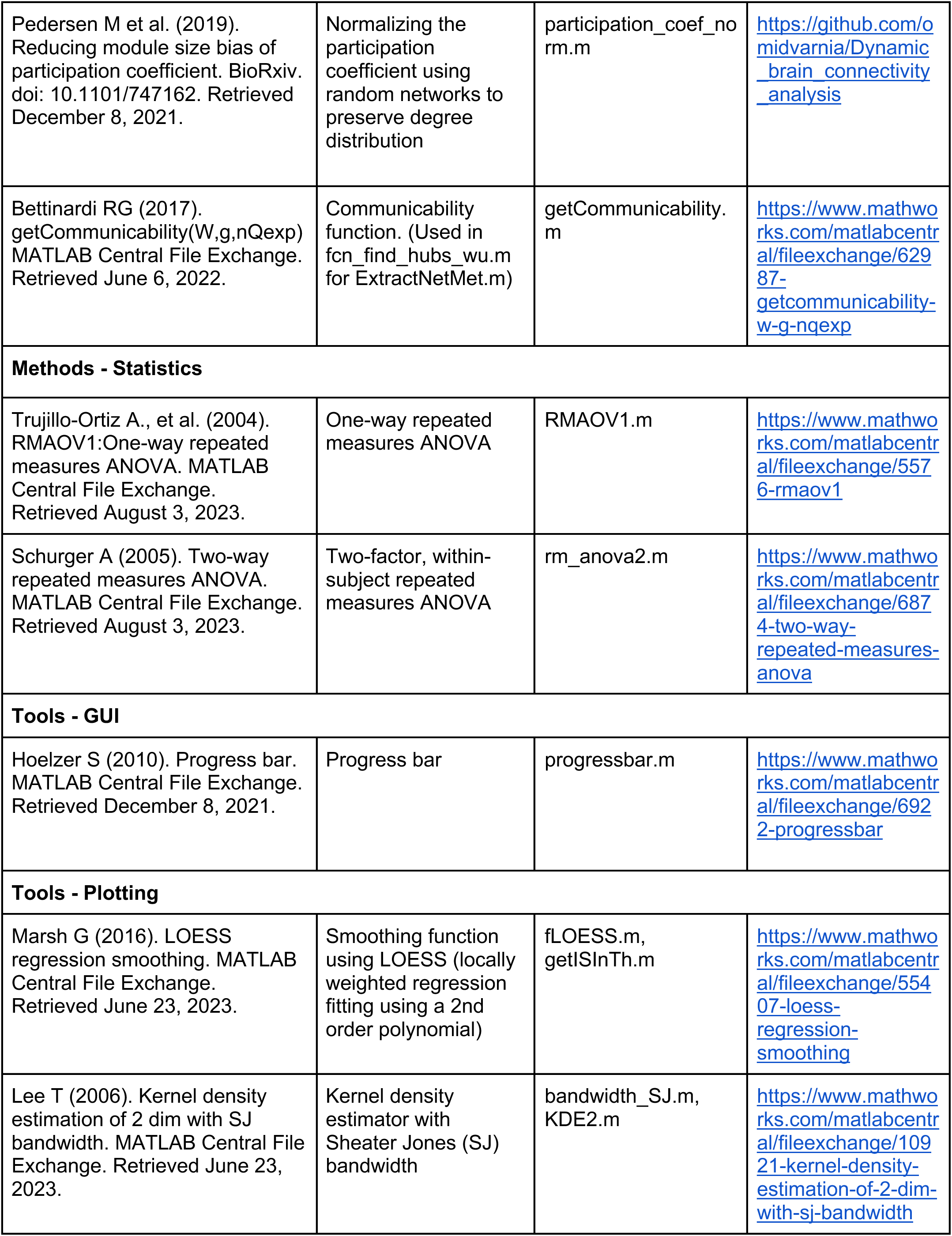
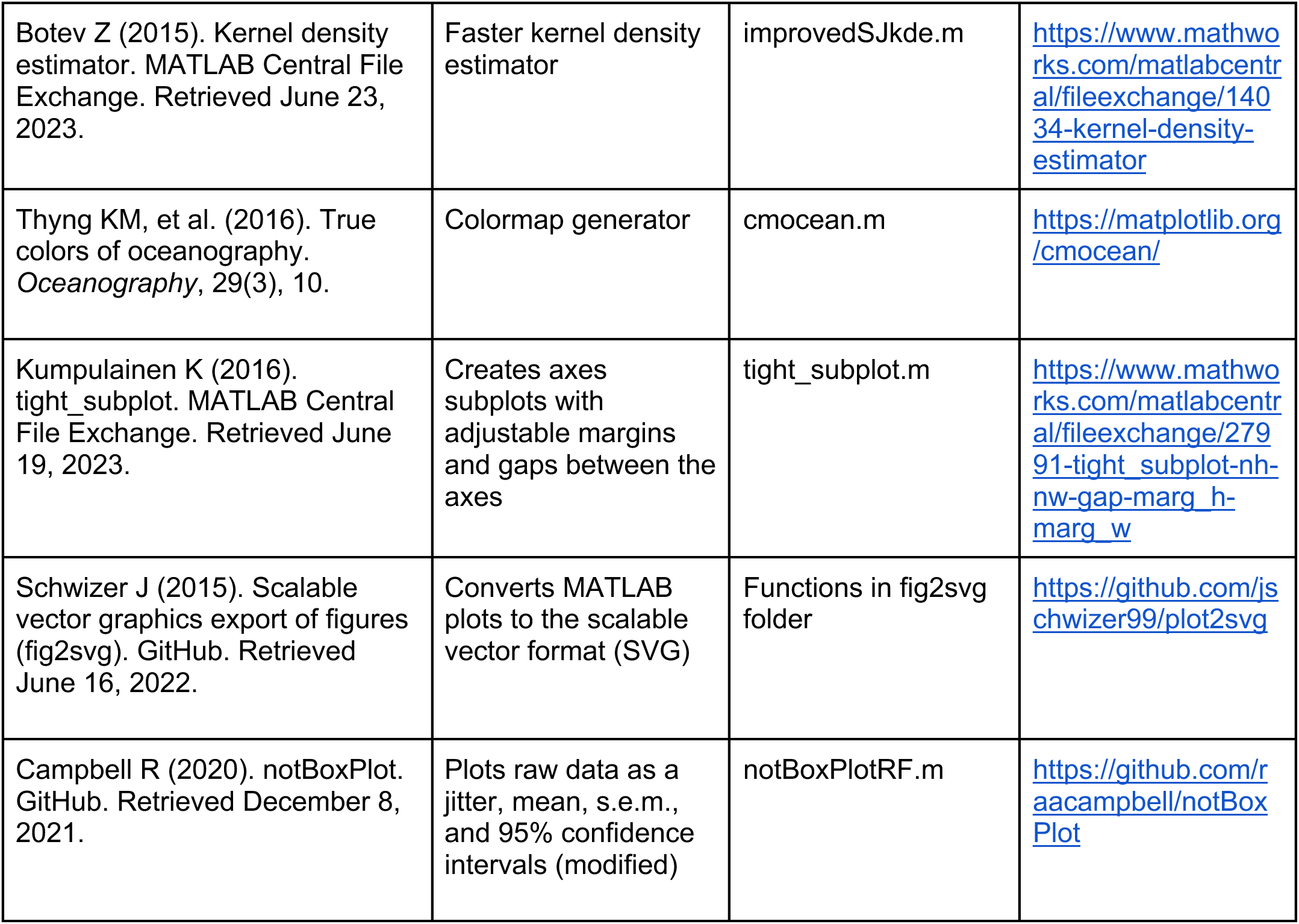
MATLAB functions or code from other sources incorporated in MEA-NAP.

**Supplemental Table S2.** Comparison with other publicly available MEA analysis or functional connectivity tools.

| Supplemental Table 2. Comparison with other publicly available MEA analysis or functional connectivity tools |  |  |  |  |  |  |  |
| --- | --- | --- | --- | --- | --- | --- | --- |
|  | Adapted for MEA data | Spike Detection | Neuronal activity comparison | Inferring functional connectivity | Network metrics | Statistical analysis | Visualization & GUI |
| MEA-NAP | ● | ● | ○ | ● | ● | ● | ○ |
| Brain Connectivity Toolbox |  |  |  | ● | ● | ● |  |
| MEA-ToolBox | ● | ○ | ○ | ○ |  |  | ○ |
| MEAnalyzer | ● | ○ | ○ | ○ | ○ |  | ○ |
| meaRtools | ● |  | ○ | ○ |  | ● |  |
| BSMART |  |  |  | ○ | ● |  |  |
| ToolConnect | ● |  |  | ○ | ● |  | ○ |
| SPICODYN | ● |  | ○ | ○ | ○ |  | ○ |
| Table Legend: Closed circle = many features; Open circle = limited features |  |  |  |  |  |  |  |

**Supplemental Table S3.**
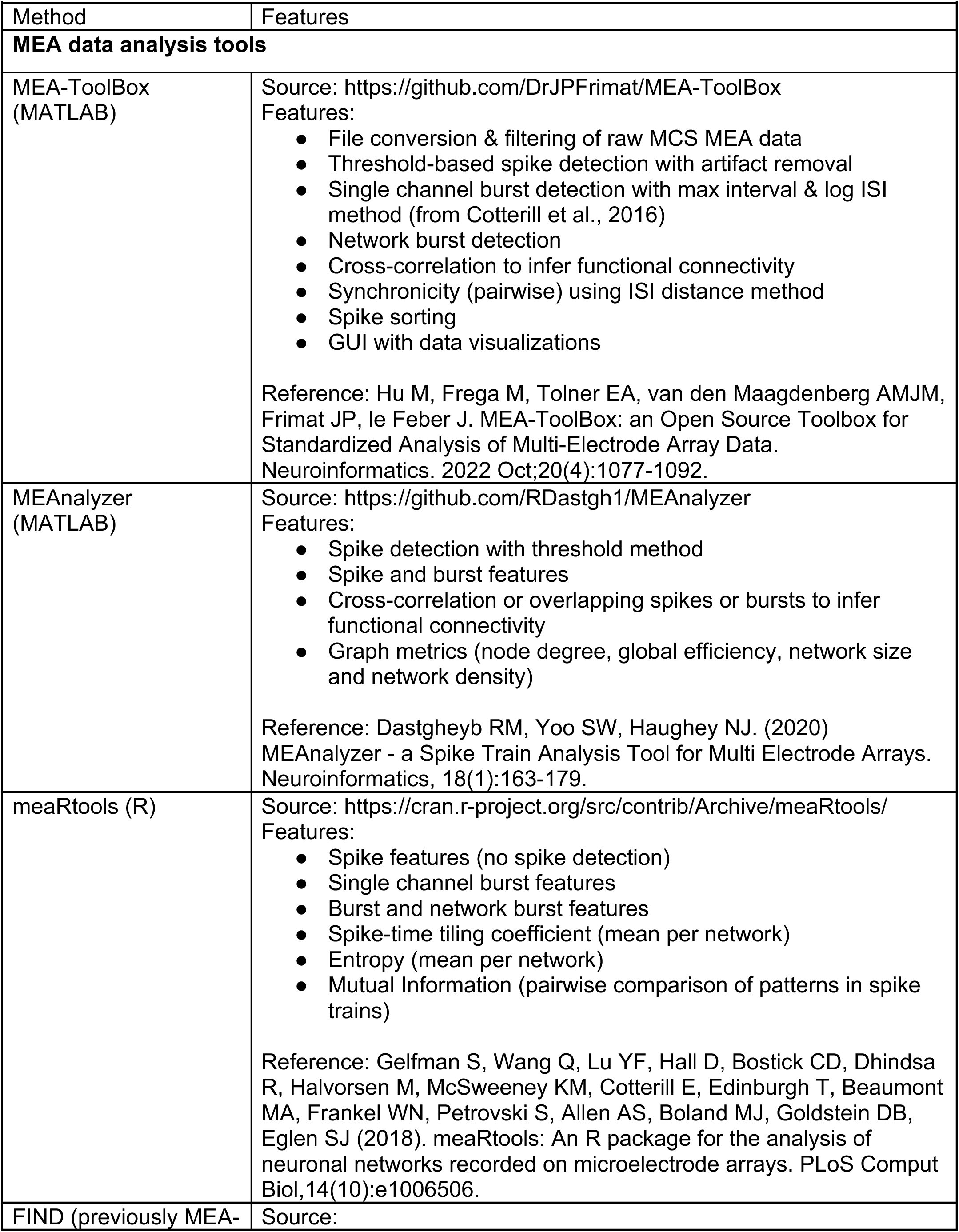
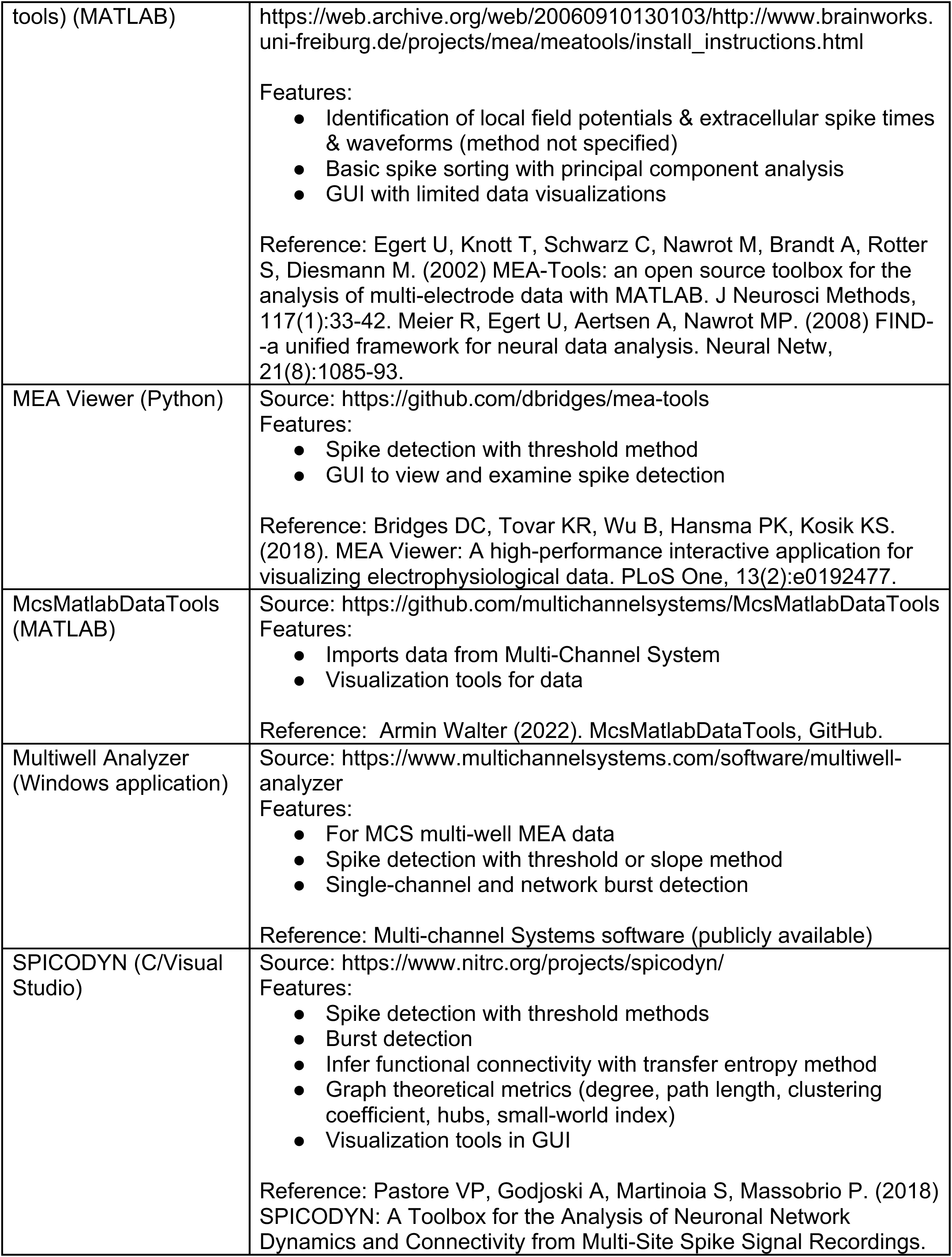
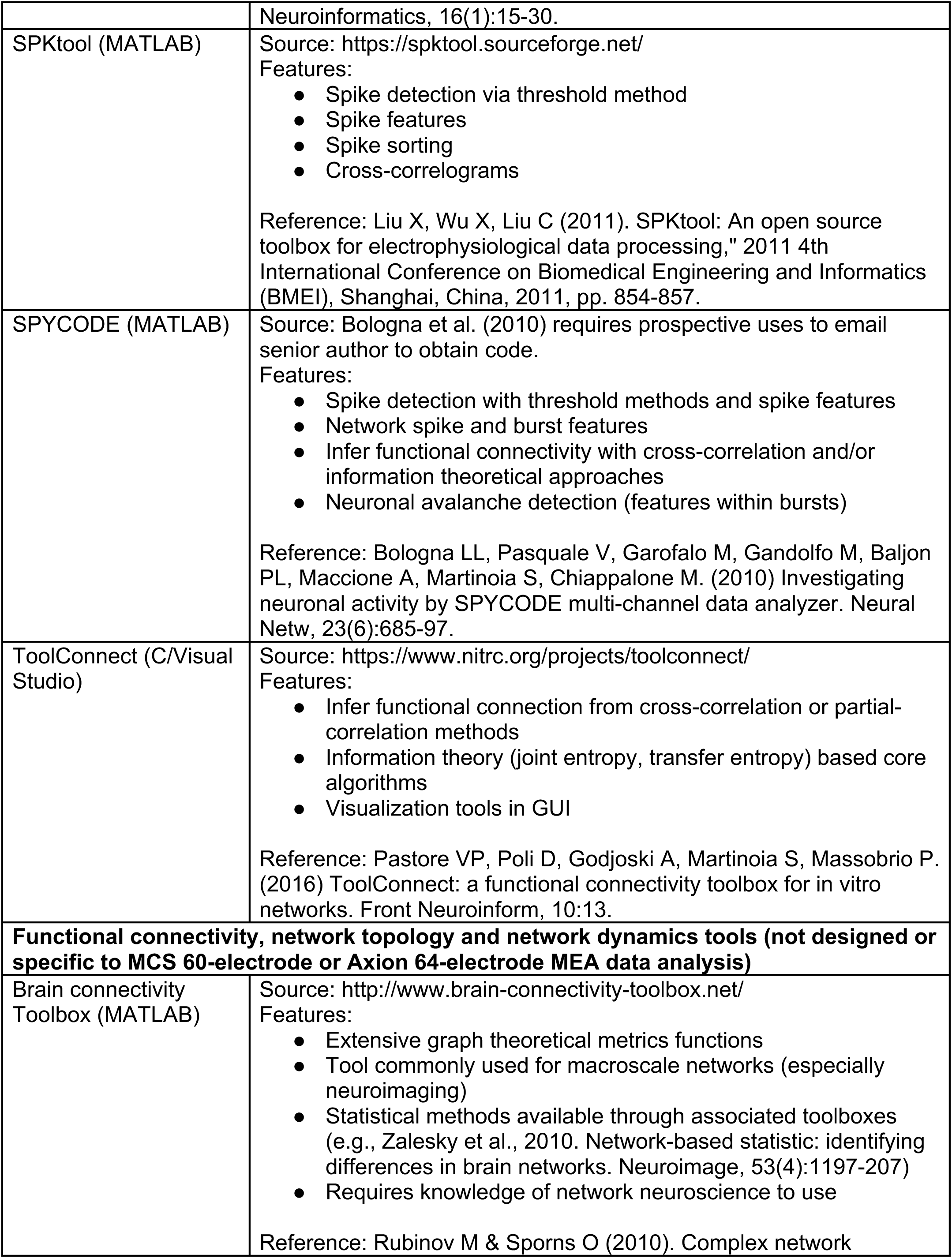
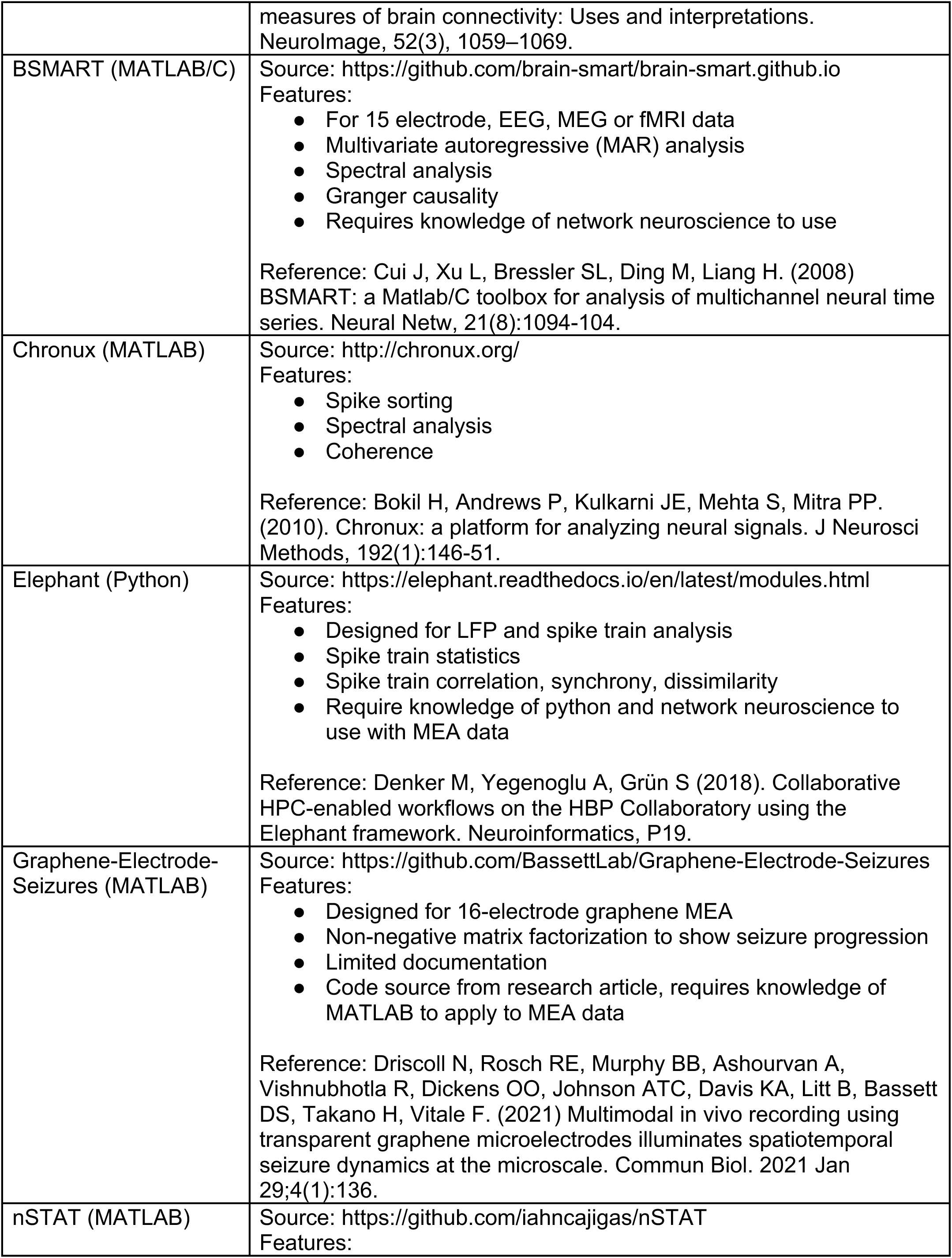
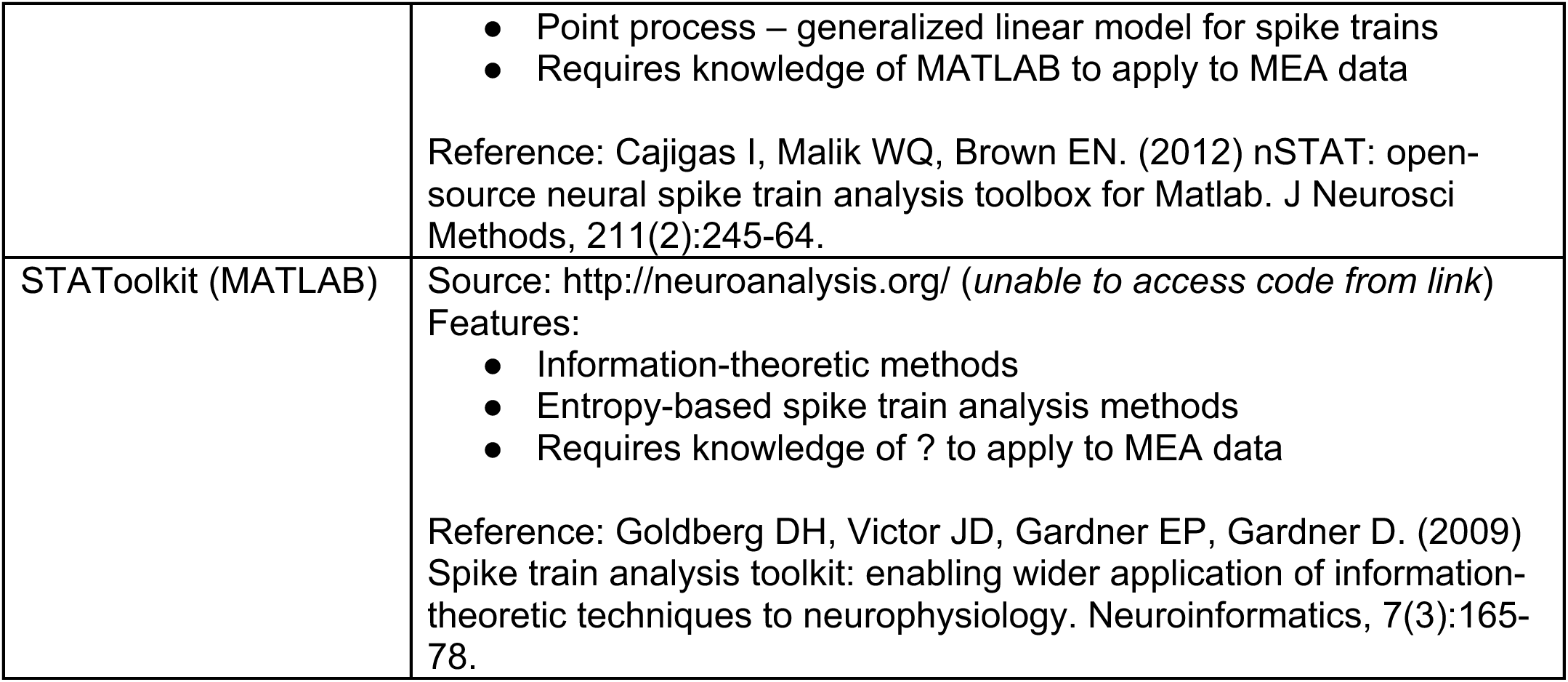
Publicly available MEA analysis or functional connectivity tools.

**Supplemental Resource 1.** Sample MEA data from 2D human iPSC-derived NGN cortical cultures. Harvard Dataverse, https://doi.org/10.7910/DVN/Z14LWA

**Supplemental Resource 2.** Sample output folder from MEA-NAP for development of 2D human iPSC-derived NGN cortical cultures. Harvard Dataverse, https://doi.org/10.7910/DVN/Z14LWA

**Supplemental Resource 3.** Video tutorial for new users of MEA-NAP. Harvard Dataverse, https://doi.org/10.7910/DVN/Z14LWA

